# An Inverse Transwell Assay for Airway Mucus Barrier Function Reveals both Virus- and Mucin-Specific Impacts on Infection

**DOI:** 10.64898/2026.07.19.739437

**Authors:** Maria Corkran, Elizabeth M. Engle, Heng Pan, Sahana Kumar, Allison Boboltz, Jeffrey R. Johnson, Gregg A. Duncan, Margaret A. Scull

**Affiliations:** Department of Cell Biology and Molecular Genetics, Maryland Pathogen Research Institute, University of Maryland, College Park, MD, USA; Molecular and Cellular Biology Program, University of Maryland, College Park, MD, USA; Fischell Department of Bioengineering, University of Maryland, College Park, MD, USA; Department of Microbiology, Icahn School of Medicine at Mount Sinai, New York, NY, USA; Global Health and Emerging Pathogens Institute, Icahn School of Medicine at Mount Sinai, New York, NY, USA

**Keywords:** mucus, mucins, lung diseases, influenza, human, rhinovirus

## Abstract

Respiratory viruses are a significant cause of morbidity and mortality world-wide and an important trigger of acute exacerbation in chronic lung disease. Secreted airway mucus – a front-line defense system against respiratory virus infection – is largely composed of glycosylated mucins that promote virus trapping via steric and adhesive interactions. Still, the degree to which mucus can trap specific viruses is unclear. Further, mucin expression is altered in chronic lung disease with undefined impacts on host susceptibility to infection. Here, we devised an inverse Transwell assay (ITA) to specifically probe the barrier function of mucus towards infection without the confounding effects of ongoing mucus secretion and transport on viral dynamics. Using the ITA, we assessed the barrier function of human airway epithelial (HAE) culture-derived mucus towards influenza (IAV), rhinovirus (RV), adenovirus, and parainfluenza virus. Results showed differences in mucus penetration efficiency between viruses, with IAV being the most inhibited relative to no mucus controls. Notably, IAV and RV penetration efficiency was similar between normal mucus and mucus sourced from an in-vitro model of asthmatic epithelium. To further explore the role of specific mucins during infection, we employed CRISPR/Cas9-modified HAE cultures lacking either MUC5B or MUC5AC expression. Direct infection in these cultures with IAV yielded higher viral titers compared to control HAE, suggesting both MUC5B and MUC5AC contribute to antiviral defense. Application of mucus harvested from specific mucin-knockout or control HAE in the ITA revealed that while RV penetration was similar across conditions, IAV was more efficient in breaching MUC5AC-depleted gels. Subsequent biophysical analysis of these mucus gels revealed a larger pore structure in the absence of MUC5AC. Together, these data indicate mucus-mediated restriction is virus dependent and highlight the contribution of MUC5AC to mucus structure and antiviral capabilities. Further, they establish the ITA as a tunable platform enabling investigation into mucus penetration by diverse viruses and the effects of altered mucus composition on barrier function.

**Clinical Relevance:** Mucus composition is altered in chronic lung disease states and during inflammation with unknown consequences on its barrier function towards respiratory virus infection. Using an inverse Transwell assay, we describe virus-specific kinetics through mucus representative of health and disease and identify a critical role for MUC5AC in defense towards influenza A virus. This work can inform future strategies to improve infection prevention or mucus targeted therapies, and may help explain differences in susceptibility to viral infections across the population.

## INTRODUCTION

Respiratory viruses cause frequent epidemics in the human population and are responsible for significant morbidity and mortality as well as economic loss. While these impacts are best exemplified by the recent COVID-19 pandemic, driven by the global spread of severe acute respiratory syndrome coronavirus 2 (SARS-CoV-2), there are numerous other clinically-relevant respiratory viruses, representing a diverse array of virus families, for which effective preventive or therapeutic treatment options are lacking.^1^ These pathogens are especially important in the context of chronic lung disease, where viral infections are a common driver of acute disease exacerbation and worsened clinical outcomes.^2,3^ Notably, innate defense mechanisms against respiratory viruses within the airways critically influence host susceptibility and infection severity both in individuals with underlying lung disease and those that are otherwise healthy. Delineating these mechanisms is essential for understanding infection outcomes and developing strategies to combat respiratory virus infections across the population.

A fundamental component of innate defense in the respiratory tract is the production and secretion of mucus which lines the conducting airways, forming a physical barrier between inhaled pathogens and the underlying epithelium. Following deposition on the airway luminal surface, viruses such as influenza A virus (IAV) and rhinovirus (RV) must breach the mucus barrier to reach target epithelial cells before mucociliary clearance mechanisms, driven by the coordinated beating of cilia on the apical cell surface, expel them from the respiratory tract. Mucus itself is a complex gel composed of water, ions, lipids, globular proteins, mucin glycoproteins, and cell debris.^4^ Among the mucins, mucin 5B (MUC5B) and mucin 5AC (MUC5AC) are essential for formation of the secreted gel and establishing its viscoelastic properties.^5–7^ Both MUC5B and MUC5AC are heavily decorated in glycans which impart a net negative charge, and harbor cysteine-rich regions at the N- and C-termini that allow for mucin oligomerization via disulfide bonds.^8^ Together, these physical properties promote the formation of a mesh-like structure that works to trap inhaled particles through both adhesive and steric interactions.

Prior work has provided further insight into how mucus restricts viral infection, revealing an important role for mucin-associated glycan-virus interactions. Here, glycans such as sialic acid (SA) within the mucus layer act as decoy receptors that can immobilize viruses like IAV and human parainfluenza virus (hPIV) that typically utilize these sugars to promote cell entry.^9^ Receptor-destroying enzymes associated with the viral particle (e.g. neuraminidase (NA) and hemagglutinin-esterase glycoproteins), however, can help to overcome this trapping mechanism and contribute to mucus penetration by catalyzing the cleavage of terminal sialic acid residues.^10–12^ Additional mucus components, such as antibodies, may also act to arrest virus mobility within the mucus gel, reducing the likelihood of infection.^13,14^ Beyond these biochemical interactions, work from our group and others has suggested that the architecture (i.e. pore size) of the mucus gel and periciliary layer plays a critical role in determining particle penetration and viral infection efficiency.^15–17^

Despite these established antiviral attributes of mucus, the mucus microenvironment is highly dynamic, and changes in the composition and physicochemical properties of secreted mucus observed during both chronic lung disease and acute infection may alter mucus-mediated defenses. Mucus hypersecretion, for example, alongside muco-obstruction, elevated mucus solids content, and alterations to mucin macromolecular forms are observed in several chronic airway disease states.^18–20^ Further, where MUC5B constitutes the majority of the mucin protein present in mucus in healthy airways, the balance of MUC5B to MUC5AC shifts substantially in asthma.^21^ Increased MUC5AC is also associated with inflammation during respiratory viral infections alongside epithelial damage and remodeling that can further alter mucin expression patterns, including changes in their terminal glycan profiles.^22–24^ These alterations may influence not only the barrier function of mucus, but also its effective transport.

Diversity across common respiratory viruses with respect to their size, shape, surface chemistry, and glycan binding properties presents further challenges in deconvoluting how mucus protects against infection and how changes in the mucus barrier may disproportionally affect susceptibility to different pathogens. Rhinoviruses (RV), for instance, are on the order of 30 nanometers in diameter, whereas filamentous virions produced during IAV or respiratory syncytial virus infections can extend to micron-scale lengths.^25,26^ Logic would suggest that these filaments may face greater steric constraints within mucus. However, evidence suggests that on the contrary, concentration of NA at the influenza virion pole drives movement towards sialic acid-rich regions and facilitates filamentous particle navigation of the mucus barrier.^10^ Together, these features highlight a central challenge: effective mucus-mediated defense depends on the mucus itself, the specific properties of the virus, and the dynamic airway environment in which the infection occurs.

Human airway epithelial (HAE) cultures grown at air-liquid interface (ALI) are able to replicate the dynamic airway microenvironment, including the establishment of a pseudostratified epithelium with multiple cell types, an extracellular mucus barrier, beating cilia and mucociliary transport of the secreted mucus. In this system, changes in mucus volume, mucus transport rate, and cilia beat frequency, all influence mucus macrorheology. Importantly, each of these factors can also influence the ability of pathogens to infect and spread within the epithelium. Therefore, while HAE cultures are invaluable for probing viral infection kinetics at the mucosal surface – something traditional 2D or monolayer tissue culture systems cannot do – their emergent properties represent confounding variables in specifically assessing the barrier function of secreted mucus towards respiratory pathogens. Additionally, studying mucus barrier function toward virus infection in HAE models of health and disease is challenging due to differences in the frequency of specific cell types, potentially impacting infection independent of mucus-mediated effects. Consequently, we sought to uncouple the barrier function of mucus from mucociliary dynamics and eliminate variation in HAE cell culture composition, to directly assess mucus-mediated protection from pathogens, and to help answer questions pertaining to the role of mucus in modulating viral infection in the airways. The inverse Transwell assay (ITA) developed here allows for the measurement of viral passage and infection through specific mucus barriers without any confounding factors, directly assessing barrier function. This approach enables analysis of how changes in mucus composition, for instance the changes observed in chronic disease, affect viral penetration through mucus, as well as comparisons between viruses without having to consider differences in cell tropism.

## RESULTS

### Establishment of an inverse Transwell assay (ITA) to assess mucus barrier function

To enable a platform capable of isolating native mucus barrier function towards a variety of viral pathogens, we envisioned a workflow in which cells grown to confluence on the basolateral side of a Transwell membrane serve as targets for infection by a virus delivered to the apical chamber. The frequency of infected cells can then be visualized by fluorescence microscopy and quantified using virus-specific CellProfiler pipelines developed for the ITA as an indication of infection efficiency (Fig. 1A). The effect of a particular mucus sample on infection can then be assessed by incorporating the mucus gel of interest into the ITA platform, in between the viral inoculum and Transwell membrane. This arrangement allows the cells to have continuous access to cell culture medium while simultaneously preventing contact between the medium and the mucus sample which is critical in order to avoid swelling of the gel and subsequent changes in total solids concentration that could affect virus dynamics.^27^ It also enables precise control of the mucus volume (barrier height) and the time given for a particular virus to penetrate the barrier; further, it supports the use of mucus gels that differ dramatically in their viscoelastic properties. To optimize the ITA set-up, we first tested the effect of Transwell membrane pore size, ranging from 0.4 to 8 microns, on the ability of Madin-Darby canine kidney (MDCK) cells to form a confluent monolayer, and thus, a uniform “sensor” to detect viruses that were able to breach the experimental mucus sample barrier. MDCK cells were seeded on the basolateral surface by inverting the Transwell membrane until the cells had adhered, then flipping the membrane back to its upright position. Cell confluency was assessed two days later by visualizing nuclei density. Notably, the two smaller pore sizes, 0.4 and 3 microns, allowed a tight monolayer to form, whereas seeding cells on the 8-micron pore size resulted in larger spaces between cells as indicated by fluorescence microscopy (Fig. 1B). Since infection of the cell monolayer in the ITA requires viruses to pass through the Transwell membrane from the apical to basolateral side, uninhibited passage of virions through the membrane is another key aspect of selecting a pore size to use in this assay. Therefore, after seeding cells on each Transwell membrane, we added A/Udorn/307/72 (H3N2), a pleomorphic influenza virus with a broad size range, to the apical surface of the membrane for 1 hour. The inoculum was then removed, and cells were later fixed and stained for viral antigen. Here we observed an increase in viral antigen positive cells on the 3- and 8-micron membranes compared to the 0.4-micron pore size membrane, suggesting that the larger pore sizes allow for more efficient IAV passage through to the cells on the basolateral side. Based on the combined cell density and infection data, we chose a Transwell membrane with 3-micron pores to use in the ITA going forward.

**Figure 1.**
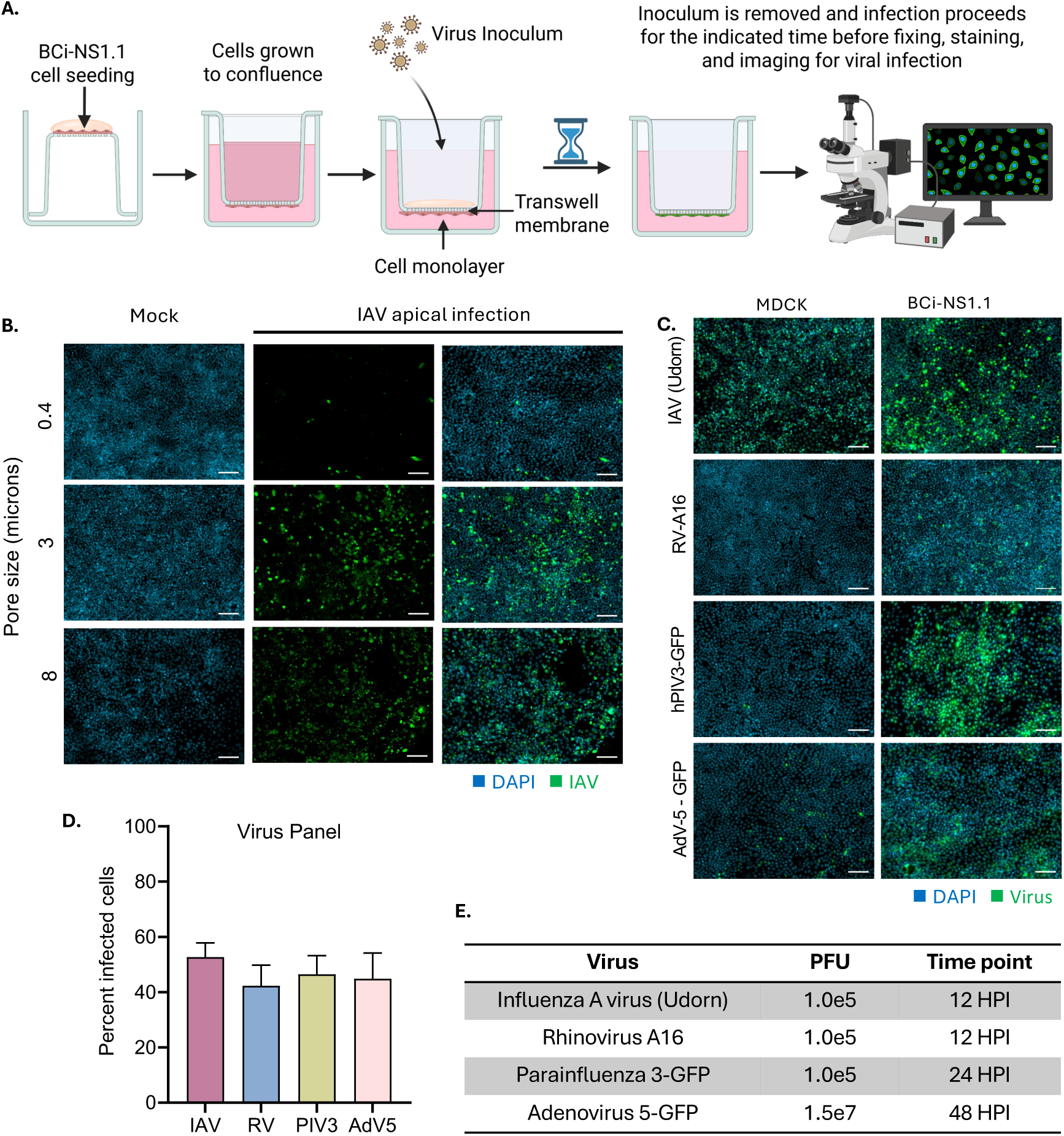
Establishment of an inverse Transwell assay (ITA) to assess mucus barrier function. (A) ITA Workflow. BCi-NS1.1 cells are seeded basolaterally on Transwell membranes before adding virus apically. Viral inoculum is removed and infection by viruses that successfully reached underlying cells is allowed to proceed. Cells are fixed, probed for viral antigen (for non-GFP-expressing viruses), and stained with DAPI. Image analysis to determine the % infected cells is done using a virus-specific CellProfiler pipeline. Created in BioRender. Scull, M. (2026) https://BioRender.com/yqunkxj. (B) Nuclei (DAPI; blue) and IAV antigen (NP; green) staining showing cell density and the frequency of infected cells, respectively, on Transwell membranes with different pore sizes. The viral inoculum was removed from the apical chamber of the Transwell after 1 hour and cells were fixed at 12 hours post infection prior to staining for viral antigen. Scale bar = 100 μm. (C) Comparison of viral infections (green) in MDCK and BCi-NS1.1 cells fixed at 12 hours post infection (IAV, RV-A16), 24 hours post infection (hPIV3-GFP), or 48 hours post infection (AdV5-GFP). Infection was visualized by staining for viral antigen or by GFP expression encoded by the viral genome. (D-E) Optimized virus dose and duration of infection for each virus selected to achieve around 50% infection. All images are representative of data collection across 2-3 independent experiments.

Our initial tests were performed with IAV, making MDCK cells, which are widely used in the field to quantify infectious IAV, a logical choice. However, since we sought to establish an assay that could be applied to different respiratory viruses, we next determined the ability of MDCK cells to support infection by a panel of diverse respiratory viruses compared to BCi-NS1.1 cells, an immortalized airway basal cell line.^28^ While IAV infection was readily detected in both cell lines, the frequency of infected cells observed for each of the other viruses, RV-A16, hPIV3-GFP, and a nonreplicating adenovirus 5 (AdV5)-GFP vector, was greater in the BCi-NS1.1 cell line than in MDCK cells given the same multiplicity of infection (MOI) and time before fixation (Fig. 1C). These data indicated that BCi-NS1.1 cells are susceptible to multiple, clinically-relevant respiratory viruses and were therefore selected for use in the ITA to ensure broad utility of our platform.

After confirming successful infection in the BCi-NS1.1 cells, we further optimized the infection parameters - including the total amount of infectious virus present in the inoculum and the duration of the infection - for each virus in our panel to achieve 50% infected cells at the time of fixation. By targeting 50% infection, the assay provides a working range that extends both above and below the baseline to visualize an increase or decrease in infection as a function of the experimental conditions. This resulted in 1×10^5^ plaque-forming units (PFU) used for each well of IAV, RV-A16, or hPIV3-GFP, allowing the infections to continue for 12 hours in the case of IAV or RV-A16, and 24 hours in the case of hPIV3-GFP (Fig. 1D, E). AdV5-GFP required a greater inoculation dose of 1.5×10^7^ PFU, and longer timeframe (48 hours) before visualization (Fig. 1D, E).

### Diverse respiratory viruses have different kinetics through healthy mucus

Following optimization of the ITA platform, we utilized our assay to interrogate the barrier function of normal mucus. To establish a reliable source of human mucus, we generated HAE cultures from normal human tracheobronchial epithelial (NHBE) cells, which, when grown under air-liquid interface conditions, express both secreted and tethered airway mucins and produce a mucus gel that can be harvested and stored for use in downstream applications, such as the ITA (Fig. 2A). To account for any donor-specific effects, we collected mucus from HAE cultures derived from three unique donors and applied these samples to the apical Transwell compartment in the ITA set-up, before overlaying the viral inoculum.

**Figure 2.**
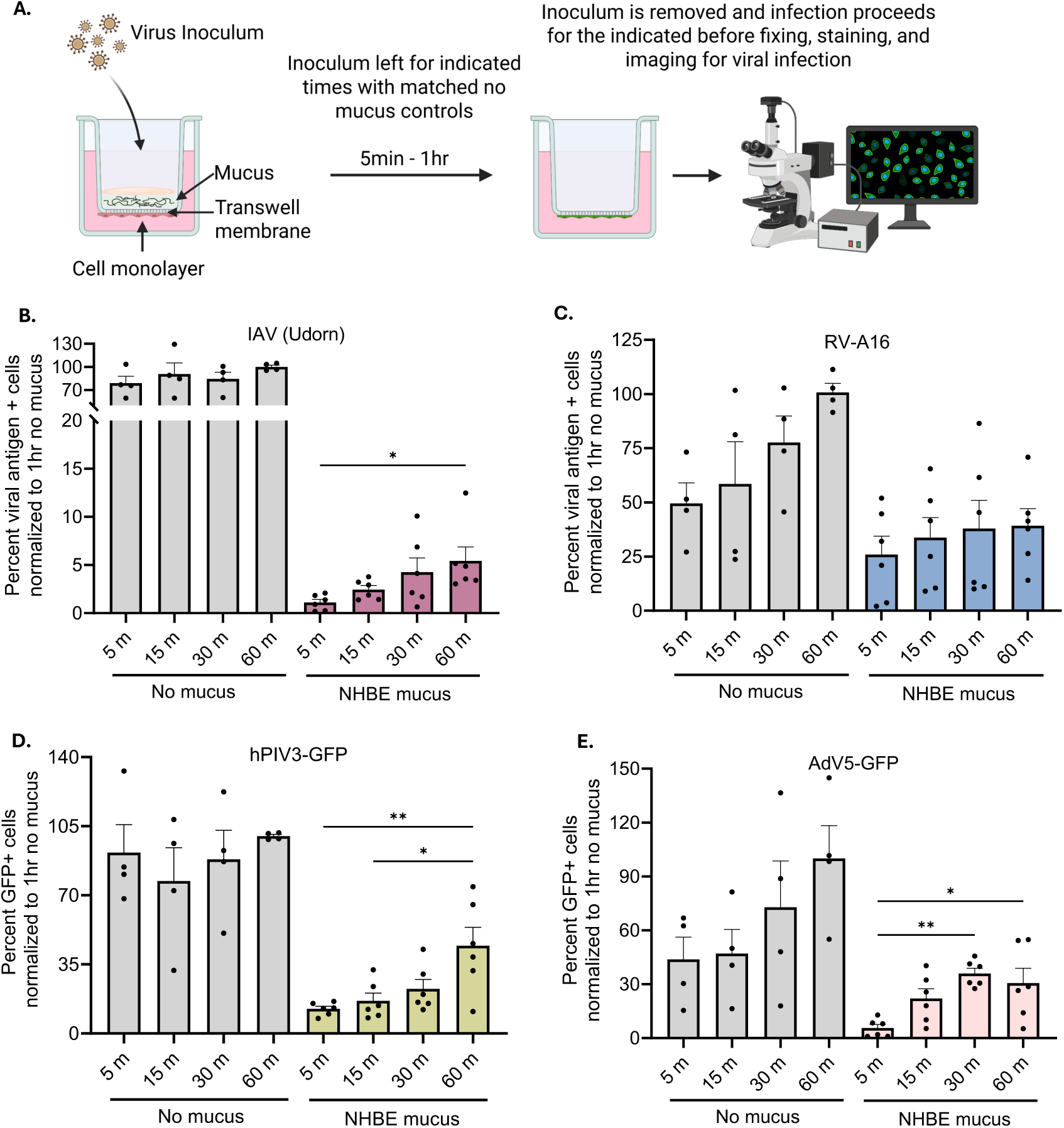
Application of the ITA to assess the barrier function of normal mucus towards diverse respiratory viruses. (A) ITA workflow. BCi-NS1.1 cells are seeded basolaterally before adding mucus into the apical Transwell chamber, followed by the viral inoculum. Virus-inoculated mucus is removed and infection by viruses that successfully penetrated the mucus barrier is allowed to proceed. Cells are fixed, probed for viral antigen (for non-GFP-expressing viruses), and stained with DAPI. Created in BioRender. Scull, M. (2026) https://BioRender.com/n7wgdf4. (B-E) IAV, RV-A16, hPIV3-GFP, and AdV5-GFP infections through normal (NHBE) mucus at indicated times compared to no mucus barrier controls. Data represent the percent viral antigen-positive cells normalized to the maximum infection condition (60 m, no mucus barrier) from n = 4-6 wells at each timepoint across 2 independent experimental replicates, analyzed via one way ANOVA with Tukey’s multiple comparisons test, *p<0.05, **p<0.01.

As expected, the ITA revealed that each virus in our panel was restricted to some extent by NHBE mucus; however, the degree of inhibition differed markedly between viruses. Among the panel of viruses, IAV displayed the greatest degree of restriction, indicating comparatively inefficient traversal of the mucus barrier that resulted in only around 5% of the number of viral antigen positive cells detected in the “no mucus” control (Fig. 2B). By comparison, inoculation with RV-A16, hPIV3-GFP, or AdV5-GFP, all resulted in at least 30% of the viral antigen positive cells detected in the corresponding “no mucus” control. Further analysis of the viral infection kinetics through mucus highlighted distinct profiles. For IAV and hPIV3-GFP, the percentage of infected cells increased gradually with longer inoculation times, consistent with a slower, time-dependent process of mucus penetration and subsequent infection (Fig. 2B, D). In contrast, we detected substantial infection achieved within the first 30 minutes for RV-A16 and AdV5-GFP with no further increase between 30 minutes and 1 hour (Fig. 2C, E). This would suggest that some fraction of the viral particles are able to navigate the mucus barrier relatively quickly, after which, when infection levels plateau, the virions remaining in the mucus may be inactivated or trapped. Together these observations suggest that while NHBE mucus acts as a barrier to each virus tested, the extent of the barrier function is virus-specific.

There are multiple factors that influence virus penetration and infection efficiency, including virion size, surface charge, and glycan binding properties, as well as the biophysical constraints imposed by the mucus mesh structure. Prior work from our group has shown that the physical structure of mucus and resulting pore sizes within mucus have a significant impact on IAV mobility.^29^ In this context, steric hindrance is a principal determinant of virion movement. As a result, virus particle size and morphology are likely to be key factors influencing virus penetration through mucus. To help understand the role of virion size in penetration and infection seen in the ITA, we performed nanoparticle tracking analysis (NTA) on each of the viruses in the panel to define their relative hydrodynamic size compared to nanoparticles of known sizes, 20 nm and 100 nm. Not surprisingly, the pleomorphic viruses IAV and hPIV3-GFP both exhibited broad size distributions, extending well beyond the 100 nm nanoparticle control measurement (Fig. S1A, S1B). In contrast, non-enveloped RV-A16 and AdV5-GFP particles that have reported sizes of 30 nm and 100 nm, respectively, displayed more uniform size profiles (Fig. S1C, S1D). The comparatively smaller average sizes of RV-A16 and AdV5-GFP could help avoid steric hindrance and facilitate more rapid diffusion of these viruses through mucus. These data are consistent with the idea that particle size affects virion penetration efficiency and subsequent infection dynamics. In addition, the NTA also measured the net surface charge of the virions in each sample, revealing a net negative charge for every virus tested (Fig. S1E). Diffusion of a negatively charged particle through a negatively charged mucus mesh could be influenced by a balance between steric hindrance and electrostatic repulsion, further altering virion transport.^30^

### Healthy and asthmatic mucus derived from in vitro HAE models have similar barrier functions against IAV or RV-A16 in the ITA

Chronic diseases of the airways, such as asthma, are complex and heterogeneous, and the underlying mechanisms that relate to infection outcomes in these individuals are not yet fully understood. Alongside chronic inflammation of the airway epithelium, a multitude of changes in mucus from asthmatic and other chronic disease states of the airways are observed including altered mucin protein secretion, increased solids content, elevated extracellular DNA and filamentous actin, and abnormal tethering of MUC5AC to the epithelial surface.^18,19,31^ Collectively, these changes can further modify the biophysical properties of mucus and consequently impact its barrier function. Still, the critical components of mucus that contribute to its barrier function, and how alterations in chronic lung disease and during infection affect susceptibility and clearance are not entirely clear.

In order to develop a hypersecretion model for asthmatic airways, HAE cultures were generated from basal cells derived from asthmatic donors (diseased human tracheobronchial epithelial cells; DHBE) and grown on Transwell inserts at ALI in parallel with normal (NHBE) donor cultures. One of the most common phenotypes observed in patients with asthma is a T2-high phenotype, or type 2-driven asthma, characterized by allergic or eosinophilic inflammation with the release of cytokines including IL-4, IL-5, and IL-13.^32^ Since prior work has established that IL-13 mediates goblet cell hyperplasia with a corresponding increase in MUC5AC secretion, the cultures derived from asthmatic donor cells were treated with IL-13.^33^ After the first 21 days of differentiation, IL-13 treatment was added to the basolateral media, with fresh IL-13 added with every media change during the final week of differentiation (Fig. 3A). Mucus from these fully differentiated cultures between days 28 and 35 was then collected and validated alongside mucus from NHBE cultures ahead of use in the ITA. As expected, IL-13 treatment increased MUC5AC secretion in NHBE cultures as well as DHBE cultures (Fig. S2). Mucus from DHBE cultures exhibited a more significant increase in total mucin content following IL-13 stimulation, resulting in mucus with the highest total mucin content, while unstimulated NHBE cultures produced mucus with the lowest total mucin content (Fig. 3B). These data suggest successful modeling of a T2-high asthmatic airway epithelium with the expected shift in mucin secretion. To evaluate barrier function, mucus collected from unstimulated NHBE-derived cultures and mucus from the stimulated DHBE-derived cultures representing the T2-high asthmatic HAE model, were selected to represent a normal baseline and a maximally altered state, respectively. Similarly, since IAV and RV differ in their basic particle composition, morphology, and glycan interactions, and because these two viruses yielded different kinetics in our initial ITA experiments (Fig. 2), they were chosen for follow up ITA experiments. Here, using these mucus samples and selected viruses in the ITA revealed the kinetics of both IAV and RV-A16 through healthy NHBE mucus are comparable to that through IL-13-stimulated DHBE mucus (Fig. 3C, D). Specifically, for IAV, viral penetration and subsequent infection through healthy mucus increased steadily from the initial 5-minute incubation through the 60-minute incubation, a trend that was similarly observed in IL-13-stimulated DHBE mucus (Fig. 3C). In contrast, RV-A16 infection through NHBE mucus, consistent with previous ITA results, showed little increase over longer incubation periods (Fig. 3D). This same pattern was seen with RV-A16 infection through IL-13-stimulated DHBE mucus. These results broadly suggest the mucus barrier derived from HAE cultures mimicking health and disease states functions similarly against either virus.

**Figure 3.**
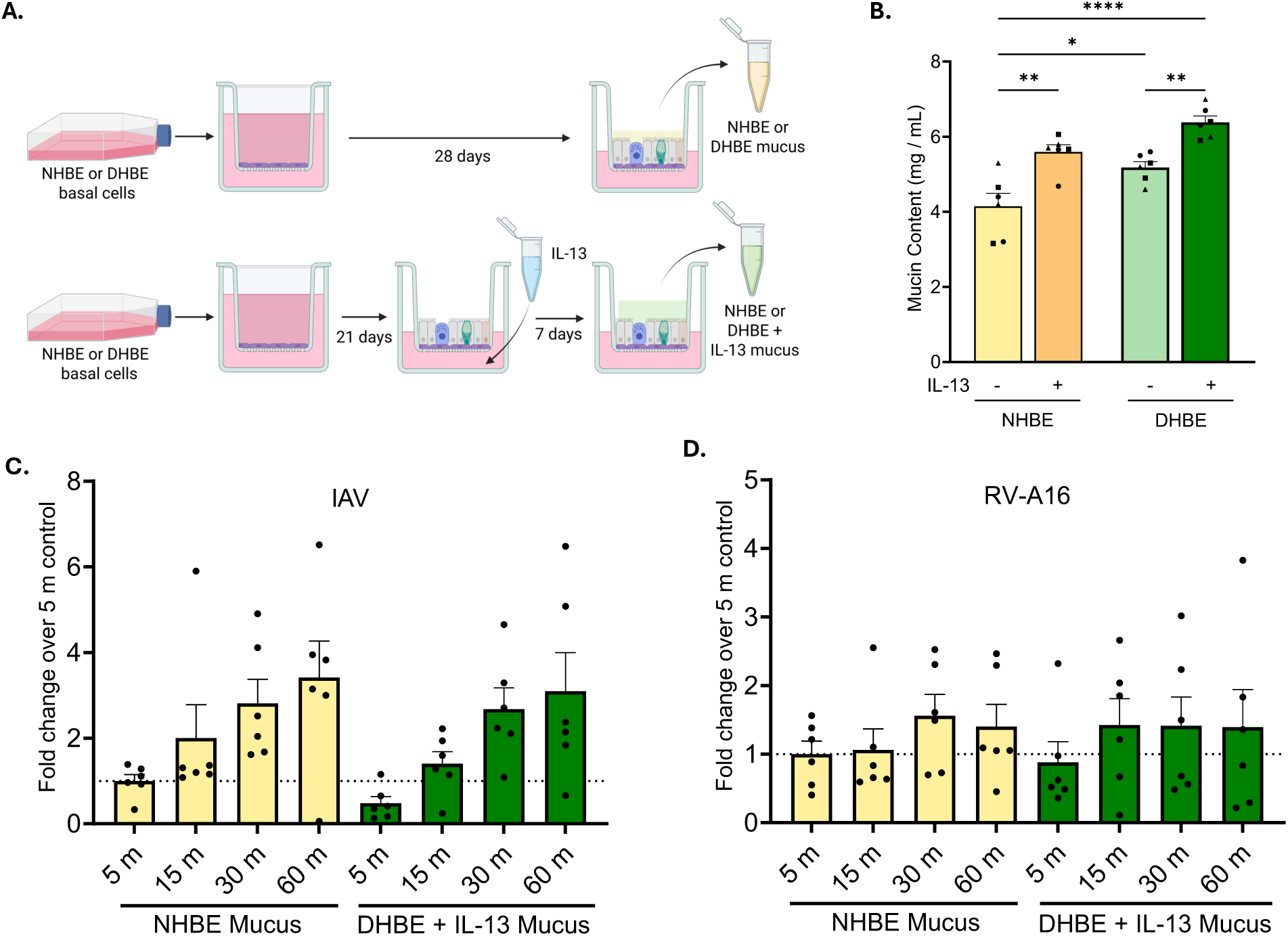
Normal and asthmatic mucus derived from NHBE and DHBE ALI cultures have similar a barrier function against IAV and RV in the ITA. (A) Schematic illustrating the NHBE and DHBE hypersecretion models. NHBE or DHBE basal cells from asthmatic donors were seeded on Transwell membranes and, once confluent, allowed to differentiate at ALI for 28 days. NHBE and DHBE HAE were stimulated with 10 ng / mL IL-13 basolaterally for the last 7 days of differentiation (bottom), or left in regular differentiation medium (top). Mucus from these cultures was collected and further analyzed for use in the ITA. Created in BioRender. Scull, M. (2026) https://BioRender.com/bc20zy1. (B) Total mucin concentration of the NHBE and DHBE cultures +/-IL-13 stimulation. Data are from n = 4-6 cultures, where individual NHBE and DHBE donors are represented by different symbols. (C-D) IAV and RV-A16 infections in the ITA using NHBE and DHBE + IL-13 mucus barriers. The percent of viral antigen positive cells is shown as the fold change over the 5 min inoculation from n = 6 well across 2 independent experimental replicates, analyzed via one way ANOVA with Tukey’s multiple comparisons test.

### IAV and RV-A16 infection in HAE models lacking MUC5B or MUC5AC expression

While the shift in the relative abundance of secreted mucin proteins, which is a major feature of chronic disease-associated mucus remodeling, did not impact virus penetration in our ITA experiments, the individual contribution of MUC5B and MUC5AC to antiviral defense remains unclear. Our group previously generated HAE cultures genetically-depleted for MUC5B or MUC5AC.^34^ Thus, we sought to utilize these models in order to determine the impact of either MUC5B or MUC5AC on viral pathogenesis. HAE knock-out (KO) cultures were infected with IAV or RV-A16 at a low MOI of 0.01 and 0.1, respectively, to allow for multiple rounds of infection, and viral titers were determined in apical washes performed on a unique set of cultures at each time point (Fig. 4A). We found that both MUC5B- and MUC5AC-KO cultures yielded elevated titers compared to those determined in control cultures expressing a non-targeting guide RNA for each virus tested. Although this did not reach significance for RV, in the case of IAV infection, there was a significant increase in viral titers later in the time course, at 48 hours (Fig. 4B, C). While these data indicate that both MUC5B and MUC5AC contribute to antiviral defense, the contribution of each mucin specifically to the barrier function towards IAV and RV-A16 is convoluted by other effects of MUC5B and MUC5AC on the mucus microenvironment. Indeed, our prior work has shown that depletion of either secreted mucin alters mucus transport.^34^ Additionally, these respiratory viruses may elicit distinct mucin expression patterns, particularly increased MUC5AC expression, after initial infection along with epithelial barrier disruption, further altering the properties of the mucus barrier.^35,36^ Since these changes could impact viral spread after initial infection, and, consequently, viral titers produced by the culture over time, the observed phenotypes across KO HAE models for both IAV and RV is difficult to fully parse.

**Figure 4.**
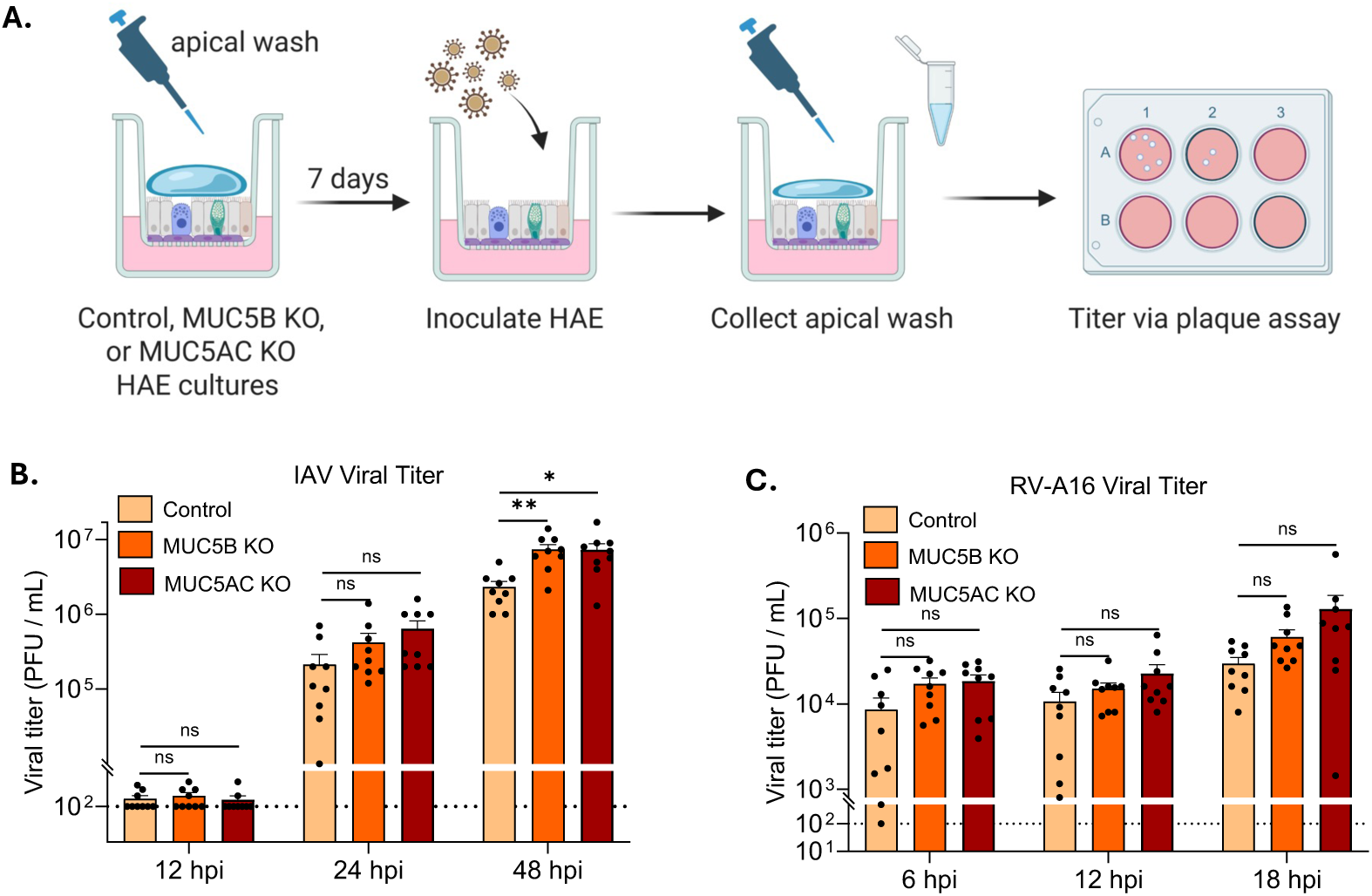
IAV and RV infection of HAE cultures genetically-depleted for MUC5B or MUC5AC. (A) Infection workflow. Mucus accumulated for 7 days prior to inoculation with IAV (A/Udorn/307/72) or RV A-16. Apical washes were collected at indicated times and viral titer determined via plaque assay. Created in BioRender. Scull, M. (2026) https://BioRender.com/fr9o0cp. (B, C) Plaque assay data shown represents n = 9 biological replicates from 3 independent experiments, analyzed via the Mann-Whitney U test, *p<0.05, **p<0.01.

### The ITA reveals distinct barrier functions of MUC5B and MUC5AC toward IAV and RV

To separate the intrinsic barrier properties of mucus from changes in mucociliary dynamics and potential differences in mucus depth across the HAE models, we harvested the mucus from differentiated MUC5B or MUC5AC KO cultures. These samples were collected in parallel with mucus from non-targeting control cultures as a physiological reference, allowing us to assess how the loss of each mucin independently affects mucus barrier function. The volume of each mucus sample was then standardized and applied to the ITA system. Following mucus application, IAV or RV-A16 was inoculated apically and allowed to penetrate the mucus barrier. Through this setup, we were able to directly evaluate the efficiency by which IAV and RV-A16 differentially traversed the mucus barriers to establish infection in underlying epithelial cells.

Our data revealed comparable RV-A16 infection across all three mucus samples when directly comparing the infection levels at each matched inoculation time, indicating that neither MUC5AC- nor MUC5B-depleted mucus differentially permits RV-A16 penetration and infection in this ITA system (Fig. 5C). Staining for viral antigen even after the longest inoculation time showed a similar number of infected cells across all mucus conditions (Fig. 5D). Similarly, for IAV, infection levels at every inoculation time through mucus depleted of MUC5B were comparable to those observed with the control mucus (Fig. 5A). This indicates that depletion of MUC5B alone also does not significantly alter IAV penetration and infection. The mucus barrier derived from the MUC5AC KO cultures, however, exhibited an increase in IAV infection over time, which was significantly higher than that of the modest increase over time seen with the MUC5B-depleted mucus barrier and the control at the longer inoculation times of 30 minutes and 1 hour (Fig. 5A). The frequency of infected cells after the 1-hour inoculation with IAV was higher through the MUC5AC-depleted mucus barrier compared to both the MUC5B-depleted and control mucus (Fig. 5B). This suggests that while the presence of MUC5AC alone may provide a baseline barrier to IAV, the significant decrease in MUC5AC may compromise the functional properties of mucus that are required for sufficient restriction of IAV. These data highlight the importance of MUC5AC to restricting IAV infection but suggest the increase in MUC5AC observed in chronic lung disease and infection does not further limit the virus, as the MUC5B-depleted mucus (i.e. MUC5AC-”rich”) has a similar barrier function to the control.

**Figure 5.**
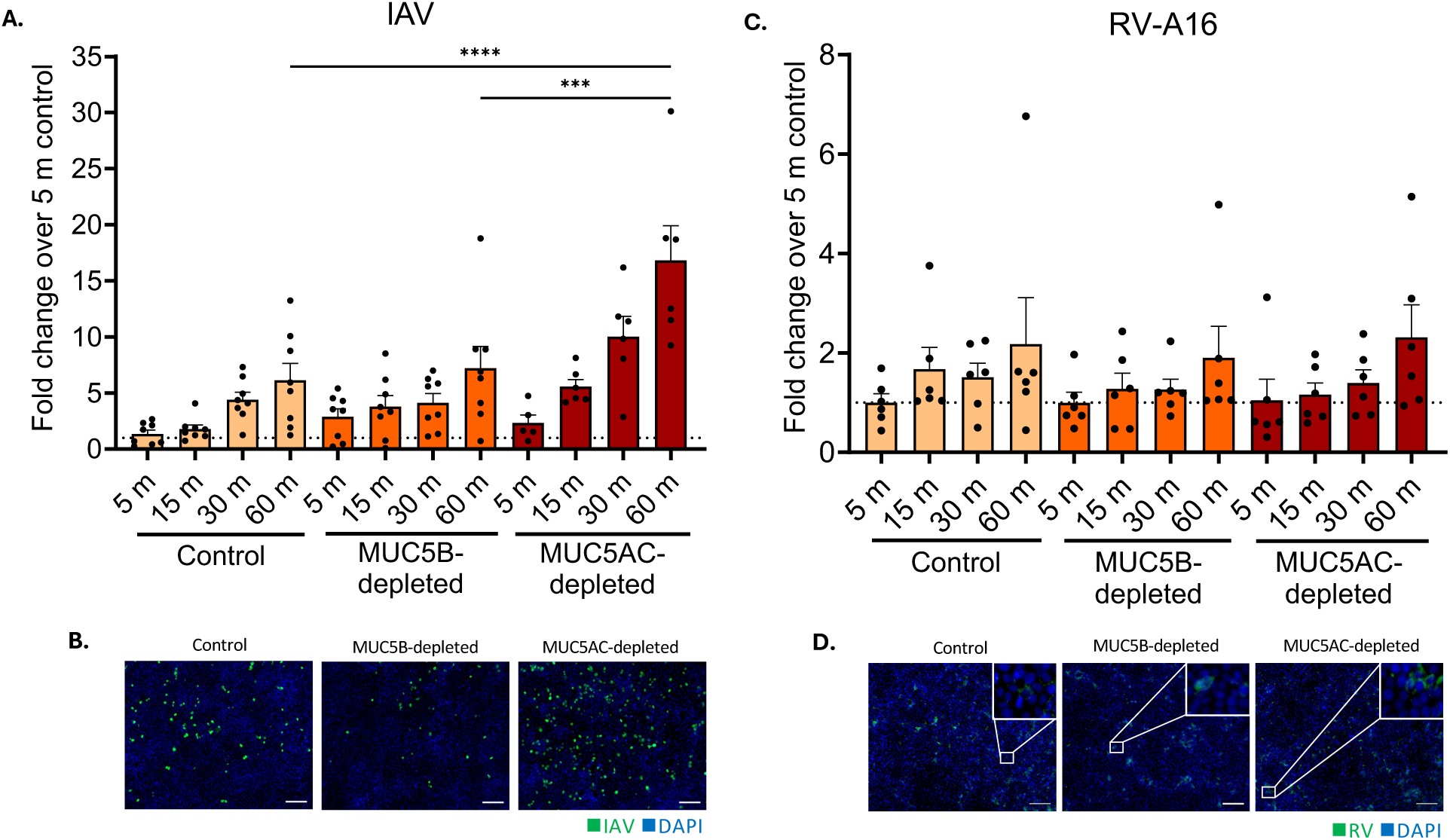
The ITA reveals distinct barrier functions of MUC5AC and MUC5B toward IAV and RV. (A, C) IAV and RV-A16 infections in the ITA using mucus from MUC5B KO and MUC5AC KO (MUC5B-depleted and MUC5AC-depleted, respectively) along with mucus from non-targeting control cultures. The percent of viral antigen positive cells is shown as the fold change over the 5 min virus inoculation through control mucus, with n ≥ 6 at each condition across 2 independent experiments, analyzed via one way ANOVA with Tukey’s multiple comparisons test, ***p<0.001, ****p<0.0001. (B, D) Representative images of IAV and RV-A16 infection (green) across each condition following a 1 h inoculation time. Scale bar = 100 µm.

### Characterization of MUC5B- and MUC5AC-depleted mucus

To delve into the underlying features of MUC5AC-depleted mucus that may contribute to its loss of barrier function towards IAV, we further characterized the mucus gels. Proteomic analysis confirmed the expected reduction in target protein expression between gels sourced from the KO and control cultures (Fig. 6A, B). While it also identified additional proteins in each mucin-depleted sample that are significantly altered compared to the control, none of these proteins to our knowledge have been ascribed pro- or antiviral properties, and are thus unlikely to impact virus traversal through mucus (Table S1). Additionally, we looked at total SA present per gram mucin protein across the samples to better understand the potential role of adhesive interactions driven by IAV hemagglutinin (HA)-glycan binding on the ability of IAV to penetrate each mucus barrier. We found that total SA was not significantly different between the mucus samples, suggesting the impact of HA binding SA is similar across conditions (Fig. 6C).

**Figure 6.**
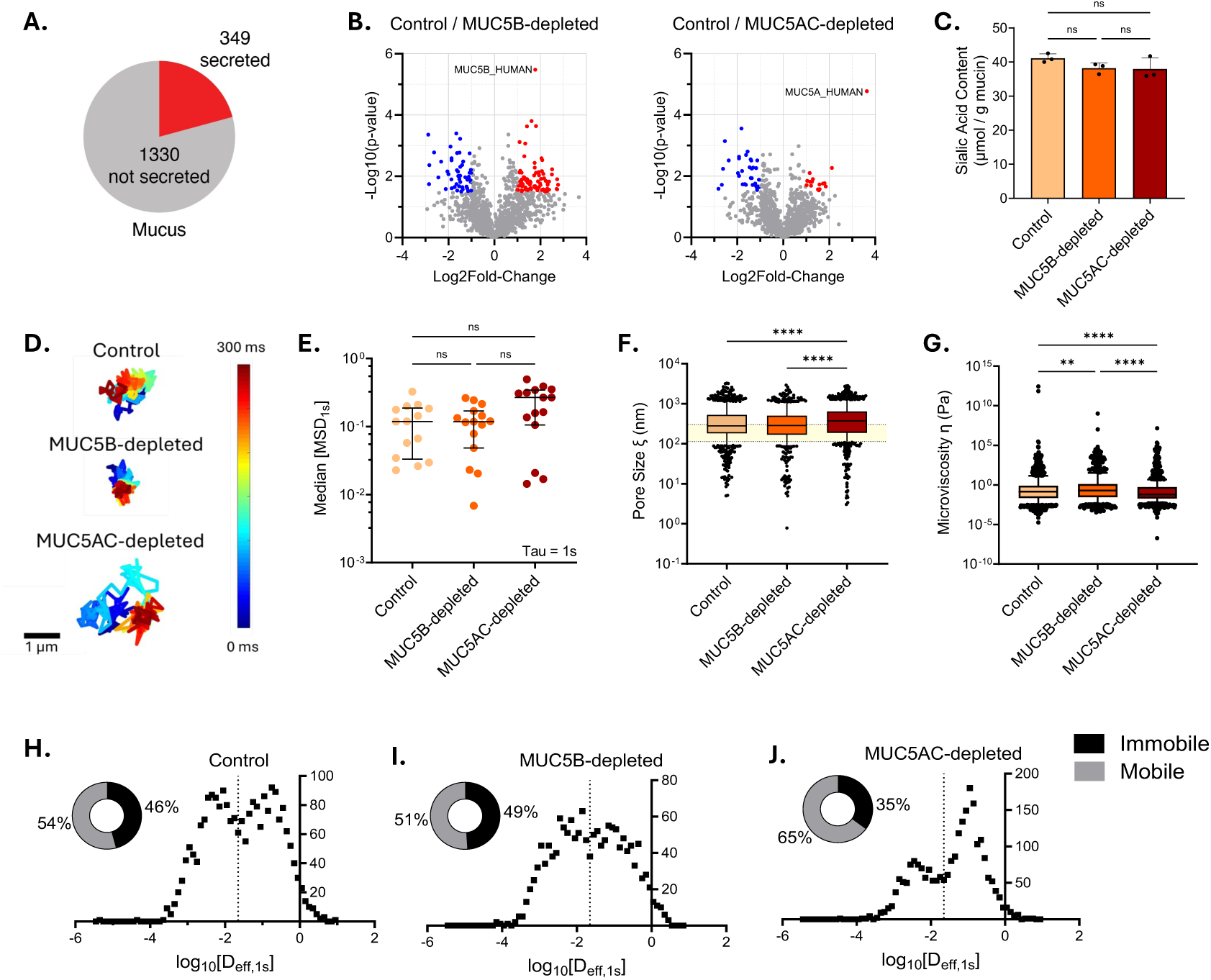
Characterization of MUC5AC- and MUC5B-depleted mucus. (A) Mass spectrometry of mucus collected from MUC5B KO, MUC5AC KO, and non-targeting control HAE, displaying total number of secreted and non-secreted proteins identified. (B) Volcano plots of proteomics data from MUC5B-depleted mucus (left) and MUC5AC-depleted mucus (right). Blue and red represent identified proteins with >1.5 -log10(p-value) and > 1 or < -1 log2Fold-Change; n = 3. (C) Sialic acid content per 1 gram mucin protein across control and knockout mucus samples; n = 3. (D) Representative trajectories of NPs within specified mucus samples. Trajectory color changes with time, with dark blue indicating 0 ms and dark red indicating 300 ms. Scale bar = 1 μm. (E) Measured mean squared displacement (MSD) at a time scale of 1 second (MSD1s) for NP diffusion in each mucus sample. Each data point represents the median measured MSD1s in each video. Black lines indicate overall median MSD1s and brackets indicate interquartile range. (F) Estimated pore size (*ξ*) from NP diffusion data. Shaded region indicates average pore size range of native mucus according to the literature.^9^ Each data point represents an estimated pore size based on the diffusion of a single NP. (G) Mucus microviscosity data extrapolated from individual MSD values of NP. (H-J) Percent fraction of mobile and immobile NPs extrapolated from individual diffusivity coefficients revealing a bimodal distribution for mobile and immobile particles.

To gain additional biophysical insights into the mucus samples, 100 nm muco-inert polystyrene nanoparticles, representative of the average hydrodynamic size of IAV, were added to each mucus sample and visualized by microscopy followed by multiple particle tracking (MPT) analysis. The diffusion rates, shown as mean squared displacement, were calculated for each tracked particle over 1 second (MSD1s) with representative trajectories shown (Fig. 6D). The resulting diffusion rates of NPs in each mucus sample, though slightly faster in the MUC5AC-depleted mucus, were not significantly different (Fig. 6E). Microrheological information extrapolated from MSD values, however, revealed that the MUC5AC-depleted mucus had larger pore sizes and lower microviscosity than MUC5B-depleted mucus and control mucus (Fig. 6F, G). The number of nanoparticles that were mobile versus immobile in each gel was then calculated based on individual diffusivity coefficients of each tracked particle, revealing a bimodal distribution representative of mobile and immobile populations. The resulting percent of particles in either population revealed a higher percent of mobile particles in the MUC5AC-depleted mucus than in the MUC5B-depleted mucus or control mucus (Fig. 6H-J). Together these data suggest that the physical restriction of IAV particles conferred by the mucus mesh of the MUC5AC-depleted mucus is less than that of the MUC5B-depleted and control mucus.

### MoNet-based predictions for NP navigation across different mucus barriers aligns with IAV infection kinetics in the ITA

Since mucus is a heterogenous gel with variable pore sizes throughout its mesh structure, the diffusivity of individual particles can vary greatly, owing to the local density of the mucins present.^37^ Further, the mobility of particles through mucus is dependent on the type of diffusion an individual particle exhibits. Through MoNet, a deep neural network, we are able to classify the NP movement of individually tracked particles, and predict the number of particles exhibiting each type of movement that are predicted to penetrate a mucus barrier of a given height in any given amount of time. As physiological mucus barriers vary greatly in their depth across different regions of the lungs, this predictive model is helpful in determining what would be expected in the ITA. Based on the diffusivities and trajectories of the tracked NPs, three diffusion modes were reviewed: Brownian motion (BM), fractional Brownian motion (FBM), and continuous time random walk (CTRW), and their survival in mucus was predicted based on previously defined machine learning and mathematical models.^27^ Based on the MoNet analysis, a majority of particles in each mucus type exhibited FBM, a subdiffusive motion (Fig. S3). Applying the diffusivity of each particle to each type of motion, we then implemented a previously-developed survival function that indicated the percent of each particle population that could navigate a defined mucus barrier, in this case one that is 10 µm thick, over a 1 hour time frame.^27^ Here we saw that a greater number of particles are predicted to cross the MUC5AC-depleted mucus barrier, and a majority of the particles that are predicted to cross exhibit BM, a random, thermally-driven motion. Overall, in alignment with the microrheological data, we observed a higher percent of NPs are mobile in MUC5AC-depleted mucus, and have a larger portion predicted to navigate across a mucus barrier in 1 hour (Fig. S3). Given the ITA results showing increased IAV infection through the MUC5AC-depleted mucus, the insight from the particle tracking analysis suggests that the greater number of larger pores present in the MUC5AC-depleted mucus, in conjunction with the decreased microviscosity, may allow for IAV to penetrate the barrier more efficiently than if MUC5AC were also present.

## DISCUSSION

This work establishes the ITA as a platform for evaluating the barrier function of airway mucus against respiratory viruses. Mucus is a critical component of airway innate defense, yet, as a complex and dynamic hydrogel, the particular aspects of mucus that enable broad protection against a diverse array of inhaled pathogens have not been fully delineated. Further, how specific changes in this barrier associated with chronic lung disease impact susceptibility to infection remain unclear. Prior studies examining the inhibitory effects of mucus toward viral infection have utilized a variety of experimental set-ups including HAE cultures at the air-liquid interface, mucus-producing cell lines such as Calu-3 cells, and purified mucins.^38–41^ While these studies have undoubtedly advanced our understanding of virus dynamics at the mucosal surface and the inhibitory properties of mucin glycoproteins, the conclusions that can be drawn regarding mucus barrier function in these models are limited. In HAE cultures, for example, infection outcomes are influenced by mucus composition as well as mucociliary activity, epithelium composition, and infection-induced epithelium remodeling. Further, neither cell line-derived mucus nor purified mucins recapitulate the biophysical properties of human airway mucus *ex vivo*.^15,42^ A key strength of the ITA described here is that it enables direct interrogation of the barrier function of any mucus gel in the absence of these confounding factors. In establishing the ITA platform, we identified Transwell membranes that support virus passage and the formation of a tight cell monolayer which is critical for maintaining assay compartmentalization and preserving the physical properties of the apical mucus layer. Our use of BCi-NS1.1 cells and optimized infection parameters also make the ITA applicable to a wide-variety of viral pathogens and allow the user to detect increases or decreases in infection across different mucus barriers.

Using the ITA to compare infection efficiency of different respiratory viruses through normal HAE-derived mucus, we found that mucus-mediated restriction is largely virus dependent. Within our virus panel, IAV was the most restricted by NHBE mucus, suggesting IAV traverses the mucus barrier less efficiently than the other viruses tested. The infection kinetics – where IAV and hPIV3-GFP exhibited a gradual rise in infection given longer inoculation times, and RV-A16 and AdV5-GFP reached maximal infection levels within the first 30 minutes – further suggest virion-specific attributes influence movement through the mucus barrier. Indeed, both glycan interactions and virus particle size likely contribute to these dynamics, as suggested by prior work.^43–46^ Mucin interactions give rise to a molecular mesh with pores typically ranging from 100 to 500 nm,^9^ or 350-900 nm as we have determined for NHBE mucus.^15^ This size range would support rapid traversal of RV-A16 and AdV5-GFP, and suggest these viruses have a high probability of reaching underlying epithelial cells in vivo before being cleared from the lung. Still, why RV and AdV5-GFP fail to achieve no mucus control-levels of infection even after longer inoculation times remains unclear, perhaps the result of particle neutralization or the formation of aggregates that are unable to reach the cell monolayer. In contrast to RV and AdV5, IAV and hPIV3-GFP infections can produce a morphologically heterogeneous population of virions that include spherical and filamentous forms.^47,48^ This heterogeneity was apparent in our NTA results and may indicate that infection kinetics are driven by morphologically-distinct pools of viral particles that move at different rates through the mucus mesh. Supporting this idea, Vahey & Fletcher demonstrated that filamentous particles exhibit a ratchet-like movement driven by the asymmetrical distribution of NA on the virion surface, unlike predictions for spherical particles.^10^ Enrichment for spherical or filamentous IAV populations by centrifugation^49^ and assay of each population separately in the ITA would provide further insights into the dynamics of these particle types in normal mucus.

Since chronic lung diseases are characterized by a multitude of changes in both the mucus microenvironment and airway epithelium, including persistent inflammation and shifts in epithelial cell type abundance, we sought to utilize the ITA to determine whether disease-associated alterations in mucus alone are sufficient to modify mucus barrier function against respiratory viruses commonly associated with disease exacerbation. We developed an in vitro asthmatic hypersecretion model, similar to those previously reported,^50,51^ in order to collect mucus representative of disease and assess its barrier function against mucus from healthy controls. Notably, when mucus from DHBE HAE stimulated with IL-13 was used in the ITA, the infection efficiency of both IAV and RV-A16 was similar to that observed in NHBE mucus. The increased mucin content in an asthmatic hypersecretion model alone is therefore not sufficient to alter virus penetration and infection. These results were unexpected given the documented increase in viral disease severity in individuals with chronic airway disease, although whether these individuals are naturally more susceptible to initial infection is less clear.^2,3^ Regardless, chronic diseases of the airway are not defined solely by mucin overproduction increasing the depth of the mucus layer, but also by changes in the secretion of extracellular DNA and actin, mucus hydration, cell shedding, and abnormal mucin tethering to the epithelium.^18,19,31^ *In vivo*, these changes may have a greater impact than the changes that occur in mucus alone, especially when coupled with the impaired mucociliary transport and epithelial remodeling that occur in disease states.

One question raised by our results comparing NHBE and DHBE+ IL-13 mucus in the ITA is whether the relative abundance of specific mucin proteins impacts mucus barrier function in more meaningful ways than total mucin content. Utilizing airway basal cells genetically-depleted for MUC5B or MUC5AC,^34^ we generated HAE models at the air-liquid interface and infected these cultures with IAV or RV-A16. While there were modest increases in viral titers across the course of infection for each virus (significant only for IAV), interpretation of these data is complicated by the impact of reduced MUC5B or MUC5AC expression on mucociliary transport dynamics. Specifically, prior data from our group indicates that the loss of MUC5B reduces mucociliary transport,^34^ and, since mucociliary transport promotes virus spread in the HAE model,^52,53^ these altered dynamics would likely result in fewer infected cells (and hence, lower viral titers) over time. However, MUC5B is also the major mucin component in the secreted mucus gel;^21,24,54^ its loss would therefore decimate the mucus barrier in the absence of compensatory mechanisms, suggesting a virus could more readily reach underlying cells to initiate infection. Since the increase in viral titers that we observed in our MUC5B- and MUC5AC-knockout HAE cultures reflects the cumulative effects of specific mucin depletion, the contribution of each mucin to barrier function is difficult to ascertain in these experiments.

This limitation is addressed by the ITA. Here, in alignment with our results in the genetically-modified HAE cultures, RV-A16 infection in the ITA was similar across conditions using mucus sourced from either specific mucin-depleted or control models. These data suggest RV is relatively “resistant” to changes in mucus composition which may explain, at least in part, why rhinoviruses are a common trigger of acute exacerbation.^3,55^ In contrast to RV, IAV infection through MUC5AC-depleted mucus was increased, suggesting impaired barrier function toward IAV in the absence of MUC5AC and highlighting respiratory virus-specific impacts of altered mucin barrier composition. While these data are consistent with prior work suggesting MUC5AC is important for defense against IAV, MUC5B-depleted gels (unlike Muc5ac-Tg mice)^56^ did not further enhance the barrier to IAV beyond that posed by mucus harvested from control cultures. The lack of congruency between studies could be due to differences in the extent of MUC5B depletion (i.e. MUC5AC enrichment) and Muc5ac upregulation between HAE and mouse models. Muc5ac upregulation in vivo was also shown to increase the thickness of the mucus barrier, whereas barrier volume is standardized in the ITA. Further, our results are consistent with earlier ITA experiments using NHBE and DHBE+IL-13 mucus where elevated levels of MUC5AC in DHBE+IL-13 mucus also had no significant impact on infection.

To probe the underlying mechanism(s) that may account for the loss of barrier function in the absence of MUC5AC, we characterized the proteome, sialic acid content, and biophysical properties of the mucus gels. Based on existing literature providing evidence for glycan-mediated trapping of IAV within human mucus^44,45,57^ and reports that glycosylation patterns differ between MUC5B and MUC5AC,^58^ we hypothesized that differences in sialic acid content between mucus gels may be responsible for the observed differences in IAV transport. However, our data indicate similar sialic acid levels across control and specific mucin-depleted gels, and instead show that the increased frequency of infected cells over time through the MUC5AC-depleted mucus in the ITA is consistent with the microrheology data. Specifically, microrheology data show that MUC5AC-depleted mucus has, on average, a greater number of larger pores and lower microviscosity than both MUC5B-depleted mucus and control. Prior work has shown unique contributions of each mucin protein to mucus architecture, where MUC5AC forms more branched polymers, yielding a mucus mesh with tighter pore structure,^59^ supporting these findings. Overall, our results are consistent with our previous work,^15^ and the idea that physical exclusion of virus particles based on pore sizes within the mucus gel is a major determinant of mucus barrier function.

While this study has established and applied the ITA to gain new insights into mucus barrier function towards respiratory virus infection, we also acknowledge several limitations in our approach. First, whether the virus-specific phenotypes observed are unique to the particular virus strains selected for our study, or are more broadly representative, remains to be seen. In addition, mucus collected from HAE culture systems is not entirely representative of the biochemical complexity of native mucus,^60^ particularly in chronic disease states where inflammatory cells can substantially alter mucus properties. Assay of mucus barrier function in the ITA is therefore ideally paired with analyses in more complex models to understand the relative contribution of differences in barrier function to infection outcomes. Importantly, the ITA has multiple applications in future work. Incorporation of airway mucus ex vivo, or mucus engineered to lack other (e.g. non-mucin) proteins, could help further define how mucus barrier function differs between individuals and which mucus components are responsible. The ITA can also be used to assess the ability of mutant viruses or those with zoonotic potential to navigate through human mucus, yielding insights into virus-mediated mechanisms of mucus penetration and species tropism. Beyond respiratory viruses, the ITA platform could also be adapted to study non-viral pathogens, engineered nanoparticles, or mucus barriers sourced from other tissues, further extending our understanding of the role that mucus plays in host defense.

## METHODS

### Cell culture

Madin-Darby canine kidney cells (MDCK; a gift from Wendy Barclay, Imperial College London), Hela H1 cells (no. CRL-1958; ATCC), and Lilly Laboratory culture - monkey kidney cells (LLC-MK2; a gift from Ralph Baric, The University of North Carolina at Chapel Hill) were maintained in high-glucose Dulbecco’s Modified Eagle’s Medium (DMEM) supplemented with 10% fetal bovine serum (FBS). MDCK and LLC-MK2 cells were 100% confluent at the time of passaging with 0.25% or 0.05% trypsin-EDTA (no. 25200-072, Gibco), respectively, and Hela H1 cells were 80-90% confluent at the time of passaging with 0.05% trypsin-EDTA. Human airway basal cells from a single donor that were previously immortalized via human telomerase reverse transcription (hTERT) and named BCi-NS1.1 cells, were generously provided by Matthew Walters and Ronald Crystal (Weill Cornell Medical College).^28^ BCi-NS1.1 cells were maintained in complete PneumaCult Ex-Plus media (no. 05040, StemCell Technologies) supplemented with 1% penicillin/streptomycin (no. 15140, Gibco) and passaged in 0.025% trypsin-EDTA at 50-60% confluence. All cells were maintained at 37°C and 5% CO2 for growth.

### Human airway epithelial (HAE) cultures

De-identified human airway (tracheobronchial) epithelial cells sourced either from individuals without underlying lung disease (normal) or donors with asthma were obtained from Lonza, Inc. (no. CC-2540S; no. 00194911S). BCi-NS1.1 cells were previously modified with CRISPR/Cas9 using guide RNAs targeting select mucin proteins (MUC5B or MUC5AC) and were sorted again in this study for GFP expression (an indication of successful transduction) to enrich for CRISPR/Cas9-edited cells prior to differentiation.^34^ Both primary and BCi-NS1.1 cells were expanded on plastic in complete PneumaCult Ex-plus media (no. 05040, StemCell Technologies) supplemented with 1% penicillin/streptomycin (no. 15140, Gibco) before seeding onto rat tail collagen type I-coated Transwell inserts (6.5 mm no. 3470, 12 mm no, 3460, Corning). Cells were seeded onto Transwells at 1×10^5^ cells / cm^2^ in complete PneumaCult Ex-Plus media and maintained under submerged conditions until confluent, at which point the apical media was removed and the basolateral media replaced with complete PneumaCult ALI media (no. 05001, StemCell Technologies) supplemented with 1% penicillin/streptomycin. Cells were maintained at 37°C and 5% CO2, and cultures differentiated for at least 28 days, with a media change 3x per week.

### HAE mucus collection

Mucus was collected from HAE cultures according to a previously detailed protocol.^61^ Briefly, fully differentiated and mucus producing HAE cultures at ALI were washed by the apical addition of 100 µL (24-well format) or 250 µL (12-well format) pre-warmed Dulbecco’s phosphate buffered saline (DPBS). Cultures were placed at 37°C for 30 min before the collection of the apical DPBS. Washes from biological replicate cultures were pooled upon collection. To remove excess DPBS, the apical washes were filtered through Amicon centrifugal filters (100 kDa, Millipore). Concentrated mucus was then collected from the filters and stored at 4°C for short term storage (<2 weeks), or -80°C for long term storage.

### Viruses

The reverse genetics system for A/Udorn/307/72 (H3N2) was a gift from Robert Lamb (Northwestern University). To amplify rescued virus, MDCK cells were grown to 100% confluence and inoculated at a low MOI of 0.001 – 0.01 in serum-free, high-glucose DMEM with the addition of 1.5 μg / mL TPCK trypsin. The infection continued for 72-96 hours, or until most of the cell monolayer had detached. The supernatant was then collected and clarified at 1,000 x *g* for 15 min at 4°C. Clarified supernatant was subsequently concentrated by ultracentrifugation through a 20% sucrose solution in NTE buffer (100 mM NaCl, 10 mM Tris pH 7.4, and 1 mM EDTA) onto a 50% sucrose cushion at 100,000 x *g* for 2 hours. The interface between the sucrose layers containing concentrated virus was collected and aliquoted for long term storage at -80°C.

Human parainfluenza virus 3 eGFP (hPIV3-GFP) was a gift from Peter Collins, Ulla Buchholz, and Cindy Luongo (NIAID). To amplify hPIV3-GFP, confluent monolayers of LLC-MK2 cells were inoculated with hPIV3-GFP at a low MOI (0.001 - 0.01) in serum-free, high-glucose DMEM with the addition of 1.5 μg/mL TPCK trypsin. The infection continued for 72-96 hours, or until most of the cell monolayer detached. The supernatant was then collected and clarified at 1,000 x *g* for 15 min at 4°C. Clarified supernatant was aliquoted for long term storage at -80°C.

Plasmid cDNA encoding the RV-A16 genome was a gift from Ann Palmenberg (University of Wisconsin-Madison). The virus was rescued by transfecting in vitro-transcribed viral RNA into HeLa H1 cells using JetPrime reagent, then further amplified at 34°C on confluent HeLa H1 cells until cytopathic effects were observed. Virus laden supernatant from the amplification was collected, clarified at 1,000 x *g* for 15 min, and frozen at -80°C prior to use.

Adenovirus 5 GFP (Ad5-Mono-EGFPΔ) was a gift from Lynda Coughlan (University of Maryland School of Medicine). This viral vector, which lacks the E1 and E3 genes, is replication deficient with EGFP as a reporter transgene and was provided as purified, high-titer aliquots.

All viruses, with the exception of AdV5-GFP, were titered according to previously developed plaque assay protocols specific for each virus: IAV,^62^ RV-A16,^63^ and hPIV3-GFP,^64^ with modifications. Briefly, cell lines corresponding to each virus amplification were seeded onto 12- or 24-well plates until fully confluent. Growth media was removed, and cells were inoculated with a small volume of 10-fold serial dilutions of the virus-containing sample in serum free DMEM for IAV and hPIV3-GFP, or McCoy’s medium supplemented with 2% FBS and 30 mM MgCl2, for RV-A16. After a 1-hour incubation, the inoculum was exchanged for a semi-solid overlay of bacteriological agar (#A5306; Sigma-Aldrich) in DMEM-F12 (no. 12500062, Gibco). For IAV and hPIV3-GFP, the overlay was supplemented with 1.5 μg / mL TPCK trypsin. Plates were then incubated at either 37°C for IAV and hPIV3-GFP, or 34°C for RV-A16 for 2-3 days until visualization of plaques by eye or with crystal violet. All virus stocks and experimental samples were only thawed once to avoid loss of infectivity.

### Virus characterization

The size and ζ-potential of viral particles were determined using nanoparticle tracking analysis (NTA; ZetaView, Particle Metrix) alongside 20 nm and 100 nm nanoparticles (NPs) as size controls. Parameters were set to a minimum brightness of 65, sensitivity of 85, frame rate of 30, and trace length of 10. For the smaller RV virions and 20 nm NPs, parameters were set to a minimum brightness of 20, sensitivity of 83, frame rate of 30, and trace length of 10.

### Inverse Transwell Array

Permeable Transwell membrane supports (6.5 mm; no. 3470, 3472, or 3464 for pore sizes 0.4, 3, and 8 µm respectively, Corning, Inc.) were inverted and coated with 30 µg / mL rat tail collagen type I (no. 354236, Corning, Inc.) in DPBS on the basolateral side of the membrane. MCDK cells or BCi-NS1.1 cells were seeded on the basolateral side of the Transwell membrane as follows: 25,000 cells in 50-75 µL of DMEM with 10% FBS (for MDCK cells) or complete PneumaCult Ex-Plus (no. 05040; StemCell Technologies) with 1% penicillin/streptomycin (no. 15140, Gibco; for BCi-NS1.1 cells) were seeded and placed (inverted) in an incubator at 37°C for 1-2 hours or until the cells were adhered to the membrane. Transwells were then removed from the incubator and another 25,000 cells in 50 µL media were added on top of the originally seeded cells and placed back in the incubator for another 1-2 hours. After that time, the Transwells were placed back upright with media added to the basolateral chamber and apical compartment, and returned to the incubator where the cells grew to 100% confluence over 1-2 days. Once at confluence, apical compartment media was removed and replaced with either a virus inoculum diluted in DPBS, or a mucus sample with virus inoculum overlay. After a set amount of time, the inoculum was aspirated off followed by a DPBS wash to remove any remaining virus that had not reached the cells. Infection by the viruses that were able to attach to cells during the given inoculation period was allowed to proceed for the indicated times. Following infection, cells were fixed in 4% paraformaldehyde in DPBS for 15 min. After fixation, cells infected with non-GFP expressing viruses were probed for viral antigen. Briefly, cells were permeabilized with 0.25% Triton-X-100, and blocked with 3% bovine serum albumin (BSA) in DPBS at room temperature for 1 hour. Primary antibodies were diluted in 1% BSA in DPBS and incubated overnight at 4°C. Samples were then washed 3 x 5 min with DPBS and secondary antibodies, similarly diluted in 1% BSA in DPBS, were added and incubated with samples for 1 hour at room temperature. Anti-IAV NP A1/A3 at 1:100 (no. MAB8251, Millipore) followed by donkey anti-mouse AF488 at 1:500 (no. A21202, Invitrogen) or anti-RV-A16 VP2 at 1:500 (no. 18758, QED) followed by goat anti-mouse IgG2b AF488 at 1:500 (no. A21141, Invitrogen) were used for IAV and RV-A16 infections, respectively. Samples were washed again 3 x 5 min with DPBS and finally stained with DAPI (#D1306, Invitrogen) for 10 min at room temperature. Samples were washed a final 3 times with DPBS before imaging. Cells infected with GFP-expressing viruses were similarly washed with DPBS and stained with DAPI before imaging. Images were obtained on a Zeiss Axio Observer 3 inverted fluorescence microscope equipped with a 10x LWD objective, Zeiss Axiocam 503 monochrome camera, and AIM-Zen 2007 software.

### Cell Profiler

Fluorescent images were exported as .TIF files and loaded onto the CellProfiler pipeline.^65^ Briefly, CellProfiler first defines nuclei and viral protein as primary objects, then relates nuclei to viral protein as “parent” and “child” objects, respectively. Related objects are further transformed into a binary relationship, assigning 0 for no relationship and 1 for parent objects labeled by child objects, identifying colocalization of cells and viral antigen. The percent of cells positive for viral antigen is directly calculated within the pipeline before being exported. As the localization of visualized viral antigen varies between viruses (IAV NP localizing to the nucleus, and RV-A16 VP2 in the cytoplasm), different CellProfiler pipelines were developed with additional steps to help CellProfiler identify and outline viral antigen-positive regions for each viral infection.

### Virus infection in HAE

Fully differentiated cultures were washed with DPBS at 37°C for 30 min to remove apical secretions. The cultures were then allowed to recover the secreted mucus layer for 7 days prior to infection. IAV and RV-A16 stocks were diluted to 500 PFU (MOI of ∼0.01) and 5,000 PFU (MOI of ∼0.1), respectively, in 10 µL DPBS for inoculation. Cultures were incubated at 37°C or 34°C for IAV and RV infections, respectively, and the virus inoculum was left on for the duration of the infection so as to not disrupt the mucus barrier. At the indicated times, washes were performed by adding 50 µL DPBS to the apical surface and incubating for 30 min at 37°C or 34°C to collect progeny virus. Washes were stored at -80°C prior to analysis via plaque assay.

### Western blotting

Mucus secretions from HAE cultures were collected in 50 µL DPBS and pooled across the same number of matched cultures. Washes were loaded with equivalent volumes in each lane of a 4-20% Tris-glycine gel (no. XP04205BOX, Invitrogen) under reducing conditions. After electrophoresis, protein was transferred to a polyvinylidene difluoride (PVDF) membrane (no. 10600030, Cytivia). The membrane was blocked in 5% fat-free milk protein in Tris-buffered saline, 0.1% Tween (TBS-T) at room temperature for 1 hour with rocking. Primary antibody (mouse anti-MUC5AC (1:1,000, no. ab3649, Abcam)) was added to 5% milk protein in TBS-T and incubated with the membrane overnight at 4°C. After incubation, the membrane was washed in TBS-T and incubated with an HRP-conjugated secondary antibody (no. sc-516102, Santa Cruz) for 1 hour at room temperature with rocking. The membrane was washed once again in TBS-T before imaging with chemiluminescent SuperSignal Dura (no. 34075, Thermo Scientific) on the iBright 1500 imaging system (Thermo Fisher).

### Particle tracking microrheology

100 nm carboxylate-modified fluorescent polystyrene nanoparticles (PS-NP) were coated with polyethylene glycol via a carboxyl-amine linkage using 5 kDa methoxy PEG-amine (no. PLS-268, Creative PEGWorks) as previously reported.^66^ The diffusion of 1 µL of the PEG coated nanoparticles (PEG-NP) in 20 µL of each mucus sample, equilibrated for 30 min at room temperature, was then measured using a Zeiss LSM 800 inverted microscope with a 63x objective as previously described.^27^ Videos were analyzed using a previously developed MATLAB code for tracking fluorescent imaged particles.^67,68^ Briefly, PEG-NPs were tracked for a total of 10 s. Mean squared displacement (MSD) was calculated for each tracked particle as 〈*MSD*(*τ*)〉 = 〈(*x*^2^ + *y*^2^)〉 where *τ* is the lag time between frames. A *τ* value of 1 s was used to minimize dynamic and static error. The generalized Stokes-Einstein relation was then used in order to find the viscoelasticity using *G*(*s*) = 2*k_B_T*/(*πas*〈Δ*r*^2^(*s*)〉) where *k_B_T* is the thermal energy, *a* is the particle radius, and *s* is the complex Laplace frequency.^66,69^ Pore size (*ξ*) and microviscosity (*η* ∗) are estimated following the complex modulus calculation, where *G* ∗ (*ω*) = *G*′(*ω*) + *G*”(*iω*) and *iω* is used in place of *s*, *i* is a complex number, and *ω* is the frequency. Then the pore size is calculated as *ξ* = (*k_B_T*/*G*′)^1/3^ and the microviscosity as *η* ∗ = (*G* ∗ (*ω*))/(*ω*).

### Machine learning analysis

For particle diffusion, a previously developed deep neural network called MotionNet (MoNet) classified the type of particle diffusion based on individual particle trajectories.^70^ This code predicts the diffusion classification, separates the trajectories by classification, and finds the anomalous diffusion exponents (α values) for each trajectory. Following this, the diffusivity for each classified particle, *D* = *MSD*/(4*τ^α^*), is used in conjunction with a previously published Mathematica code to determine the predicted particle traversal times through mucus, converted to a percentage of particles predicted to cross in 60 minutes.^71^

### Sialic acid content

Total sialic acid was quantified using a sialic acid assay kit (no. MAK314, Sigma Aldrich) according to the manufacturer’s protocols. Briefly, 20 µL mucus samples were hydrolyzed to release bound sialic acid. Total sialic acid was then oxidized, reacting with thiobarbituric acid for a colorimetric readout. The samples were read at 549 nm alongside a standard curve of known sialic acid concentration to determine total sialic acid concentration of each sample using a Multiskan Ascent microplate reader (no. 354, Labsystems).

### Mucin content

Mucin concentration was determined by alkaline 2-cyanoacetamide (CNA) fluorometry, based on a previously established protocol for O-linked glycoproteins.^72^ Briefly, mucus samples were incubated with alkaline CNA reagent containing 0.15 M NaOH at 100°C for 30 min, followed by the addition of 0.6 M borate buffer at a pH of 8.0. The resulting fluorescence intensity was measured at 336 nm excitation and 383 nm emission using the Spark Multimode microplate reader (Tecan). The mucin content was determined relative to a standard curve generated using purified mucin standards.

### Sample preparation for proteomics analyses

Samples were incubated in a buffer containing 2 M urea, 50 mM Tris-HCl, pH 8.0, and 1 mM DTT at 37 °C for 30 min. Iodoacetamide was then added to a final concentration of 3 mM and samples were incubated at room temperature for 45 min in the dark. DTT was added to a final concentration of 3 mM and 750 ng of trypsin (trypsin gold, Promega) was added to each sample, followed by overnight incubation at 37 °C with shaking. The supernatant was transferred to a fresh tube and an additional 500 ng trypsin added to the supernatant, followed by incubation for 2 h at 37 °C. Digested samples were desalted on BioPureSPN Mini C18 SPE columns (Nest Group) according to the manufacturer’s protocol. Samples were dried by vacuum centrifugation and resuspended in 0.1% formic acid (FA) for mass spectrometry analysis.

### Mass spectrometry data acquisition

All samples were analyzed on an Orbitrap Eclipse mass spectrometry system equipped with an Easy nLC 1200 ultra-high pressure liquid chromatography system interfaced via a Nanospray Flex nanoelectrospray source (Thermo Fisher Scientific). Samples were injected onto a fritted fused silica capillary (30 cm × 75 μm inner diameter with a 15 μm tip, CoAnn Technologies) packed with ReprosilPur C18-AQ 1.9 μm particles (Dr. Maisch GmbH). Buffer A consisted of 0.1% Formic Acid (FA), and buffer B consisted of 0.1% FA/80% Acetonitrile (ACN). Peptides were separated by an organic gradient from 5% to 35% mobile buffer B over 120 min followed by an increase to 100% B over 10 min at a flow rate of 300 nL / min. Analytical columns were equilibrated with 3 μL of buffer A.

The mass spectrometer acquired data in a data-independent analysis (DIA) manner. A full scan was collected at 60,000 resolving power over a scan range of 390-1010 *m/z*, an instrument controlled AGC target, an RF lens setting of 30%, and an instrument controlled maximum injection time, followed by DIA scans using 8 m/z isolation windows over 400-1000 *m/z* at a normalized HCD collision energy of 28%.

The DIA-NN algorithm was used to identify peptides/proteins and to extract intensity information from DIA data.^73^ Data were searched against the *Homo sapiens* reference proteome sequences in the UniProt database (one protein sequence per gene, downloaded on August 23, 2023). Search parameters included a fixed modification for carbamidomethyl cysteine and variable modifications for N-terminal protein acetylation and methionine oxidation. All other search parameters were DIA-NN factory defaults.

Statistical analysis of proteomics data was conducted utilizing the MSstats package in R.^74^ All data were normalized by equalizing median intensities, the summary method was Tukey’s median polish, and the maximum quantile for deciding censored missing values was 0.999. For protein abundance analyses, only the top 20 peptide features per protein were considered.

### Statistical analysis

Statistical analysis was performed using GraphPad Prism Software version 11.0.0 (84) for Windows. Additional details are provided in the figure legends.

## Author contributions

Conceptualization: M.C., G.A.D., and M.A.S; Methodology: M.C, E.M.E., J.R.J., G.A.D., and M.A.S.; Investigation: M.C., E.M.E., H.P., S.K., and A.B.; Visualization: M.C. and E.M.E; Supervision: J.R.J, G.A.D, and M.A.S.; Writing-original draft: M.C. and M.A.S.; Writing-reviewing & editing: All authors.

## Acknowledgements

We thank Hannah Zierden and Zierden lab members for providing access to their ZetaView instrument and for assistance with nanoparticle tracking analysis. We acknowledge the BioWorkshop core facility in the Fischell Department of Bioengineering at the University of Maryland – College Park for use of their Tecan Spark Multimode Microplate Reader and BD FACS Melody Cell Sorter and are also grateful to the directors and teams of the MPRI Flow Cytometry and Cell Sorting Facility.

