## Supplemental Figures for "An Inverse Transwell Assay for Airway Mucus Barrier Function Reveals both Virus- and Mucin-Specific Impacts on Infection"

### SUPPLEMENTAL FIGURE LEGENDS

**Figure S1. Virus particle size distribution and net surface charge.** (A-D) Measured hydrodynamic size distribution of each virus preparation in comparison to nanoparticle size controls via NTA. Dotted lines indicate the range in which 10% to 90% of measured particles fall. The dashed line indicates the measured median particle size for each virus sample. (E) Net surface charge (mV) of virus particles measured via NTA.

**Figure S2. IL-13 treatment leads to increased MUC5AC expression across NHBE and DHBE cultures.** Western blot of representative mucus samples derived from NHBE and DHBE cultures with and without IL-13 treatment. Mucus was collected by washing the apical culture surface with DPBS and pooled across 3 individual cultures for each condition. Culture washes were separated by electrophoresis under reducing conditions and probed for MUC5AC.

**Figure S3. Machine learning analysis of NP movement through MUC5AC- and MUC5B-depleted mucus.** (A) Fraction of mobile NPs exhibiting each type of motion across mucus samples. (B) Survivability of NPs as predicted to cross a mucus barrier of 10 μm depth over 1 hour. (C) Representative trajectories of mobile NPs exhibiting each type of motion.

**Table S1. Proteomics Data Summary.** Summary of analysis (Tab 1) and list of proteins identified with Log2FC of either $\geq$ 1 or $\leq$ -1 for control / MUC5AC-depleted (Tab 2) or control / MUC5B-depleted (Tab 3).

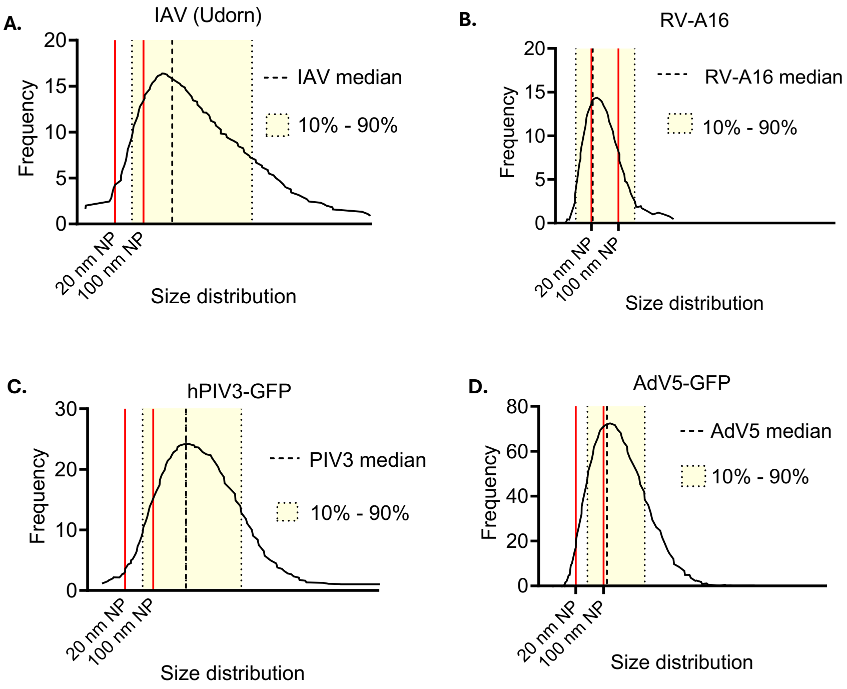


E.

| | IAV (Udorn) | RV-A16 | hPIV3-GFP | AdV5-GFP |
|---|---|---|---|---|
| Net surface charge (mV) | -34.26 ± 0.9 | -6.95 ± 1.07 | -29.06 ± 1.56 | -11.02 ± 1.67 |

**Figure S1.**

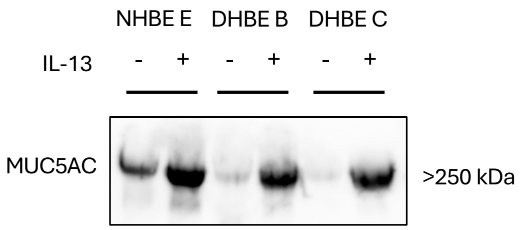


**Figure S2.**

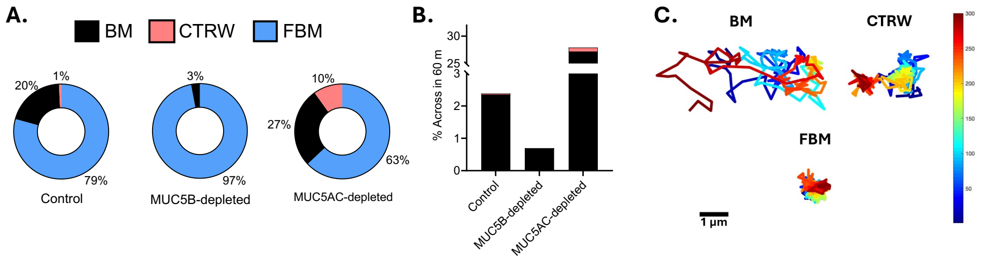
A.
BM
CTRW
FBM
20%
1%
79%
Control
3%
97%
MUC5B-depleted
10%
27%
63%
MUC5AC-depleted
B.
% Across in 60 m
30
25
3
2
1
0
Control
MUC5B-depleted
MUC5AC-depleted
C.
BM
CTRW
FBM
1 µm

**Figure S3.**

Table S1.

| WT-MUC5AC KO | |
|---|---|
| **Column Name** | **Description** |
| Protein | UniProt protein accession(s). Accessions separated by semi-colons represent identifications by non-unique peptide sequences mapping to multiple proteins. |
| Protein.Description | UniProt protein description (FASTA header line) |
| Protein.Name | UniProt unique entry name |
| WT-MUC5ACKO_log2FC | Log2fold-change comparing non-targeting control and MUC5AC KO conditions, organized by decreasing fold change values, >1 or <-1. |
| NEG LOG 10 (PVALUE) | Negative log10 transformation of computed p-value following Msstats-computed p-value comparing intensities comparing conditions indicated |

| WT-MUC5B KO | |
|---|---|
| **Column Name** | **Description** |
| Protein | UniProt protein accession(s). Accessions separated by semi-colons represent identifications by non-unique peptide sequences mapping to multiple proteins. |
| Protein.Description | UniProt protein description (FASTA header line) |
| Protein.Name | UniProt unique entry name |
| WT-MUC5BKO_log2FC | Log2fold-change comparing non-targeting control and MUC5B KO conditions, organized by decreasing fold change values, >1 or <-1. |
| NEG LOG 10 (PVALUE) | Negative log10 transformation of computed p-value following Msstats-computed p-value comparing intensities comparing conditions indicated |

| | A | B | C | D | E |
|---|---|---|---|---|---|
| 1 | Protein | Protein.Description | Protein.Name | WT-MUC5ACKO_log2F | NEG LOG 10 (PVALUE) |
| 2 | Q92888 | ARHG1_HUMAN Rho guanine nucleotide exchange factor 1 OS=Homo sapiens OX=9606 GN=ARHGEF1 PE=1 SV=2 | ARHG1_HUMAN | 1.83598425 | 1.40195417 |
| 3 | P0C672 | TSN19_HUMAN Tetraspanin-19 OS=Homo sapiens OX=9606 GN=TSPAN19 PE=3 SV=1 | TSN19_HUMAN | 1.771828655 | 1.781294794 |
| 4 | O75190 | DNJB6_HUMAN DnaJ homolog subfamily B member 6 OS=Homo sapiens OX=9606 GN=DNAJB6 PE=1 SV=2 | DNJB6_HUMAN | 1.447440043 | 2.402080451 |
| 5 | P08263 | GSTA1_HUMAN Glutathione S-transferase A1 OS=Homo sapiens OX=9606 GN=GSTA1 PE=1 SV=3 | GSTA1_HUMAN | 1.412170392 | 1.979195352 |
| 6 | Q9Y4F9 | RIPR2_HUMAN Rho family-interacting cell polarization regulator 2 OS=Homo sapiens OX=9606 GN=RIPOR2 PE=1 SV=4 | RIPR2_HUMAN | 1.398191113 | 2.313714373 |
| 7 | P05155 | IC1_HUMAN Plasma protease C1 inhibitor OS=Homo sapiens OX=9606 GN=SERPING1 PE=1 SV=2 | IC1_HUMAN | 1.364800526 | 1.874428707 |
| 8 | P07711;Q5NE16 | CATL1_HUMAN Procathepsin L OS=Homo sapiens OX=9606 GN=CTSL PE=1 SV=2;CATL3_HUMAN Putative inactive cathepsin L-like protein CTSL3P OS=Homo sapiens OX=9606 GN=CTSL3P PE=5 SV=1 | CATL1_HUMAN;CATL3_HUMAN | 1.157931667 | 1.762097495 |
| 9 | O43854 | EDIL3_HUMAN EGF-like repeat and discoidin I-like domain-containing protein 3 OS=Homo sapiens OX=9606 GN=EDIL3 PE=1 SV=1 | EDIL3_HUMAN | 1.14916606 | 1.56561681 |
| 10 | P01133 | EGF_HUMAN Pro-epidermal growth factor OS=Homo sapiens OX=9606 GN=EGF PE=1 SV=2 | EGF_HUMAN | 1.13685733 | 1.515537283 |
| 11 | P11142;P0DMV9;P0DMV8 | HSP7C_HUMAN Heat shock cognate 71 kDa protein OS=Homo sapiens OX=9606 GN=HSPA8 PE=1 SV=1;HS71B_HUMAN Heat shock 70 kDa protein 1B OS=Homo sapiens OX=9606 GN=HSPA1B PE=1 SV=1;HS71A_HUMAN Heat shock 70 kDa protein 1A OS=Homo sapiens OX=9606 GN=HSPA | HSP7C_HUMAN;HS71B_HUMAN;HS71A_HUMAN | 1.121526059 | 2.02564814 |
| 12 | Q01344 | IL5RA_HUMAN Interleukin-5 receptor subunit alpha OS=Homo sapiens OX=9606 GN=IL5RA PE=1 SV=2 | IL5RA_HUMAN | 1.067651978 | 1.549095672 |
| 13 | Q9BYE2;P07477 | TMPSD_HUMAN Transmembrane protease serine 13 OS=Homo sapiens OX=9606 GN=TMPRSS13 PE=2 SV=5;TRY1_HUMAN Serine protease 1 OS=Homo sapiens OX=9606 GN=PRSS1 PE=1 SV=1 | TMPSD_HUMAN;TRY1_HUMAN | 1.010501021 | 1.55938818 |
| 14 | P17405 | ASM_HUMAN Sphingomyelin phosphodiesterase OS=Homo sapiens OX=9606 GN=SMPD1 PE=1 SV=5 | ASM_HUMAN | -1.014744691 | 1.806055783 |
| 15 | Q14315;O75369 | FLNC_HUMAN Filamin-C OS=Homo sapiens OX=9606 GN=FLNC PE=1 SV=3;FLNB_HUMAN Filamin-B OS=Homo sapiens OX=9606 GN=FLNB PE=1 SV=2 | FLNC_HUMAN;FLNB_HUMAN | -1.027907555 | 2.928779821 |
| 16 | O43745 | CHP2_HUMAN Calcineurin B homologous protein 2 OS=Homo sapiens OX=9606 GN=CHP2 PE=1 SV=3 | CHP2_HUMAN | -1.035444234 | 1.466440836 |
| 17 | O60613 | SEP15_HUMAN Selenoprotein F OS=Homo sapiens OX=9606 GN=SELENOF PE=1 SV=4 | SEP15_HUMAN | -1.051010429 | 1.910355478 |
| 18 | Q9UKR3 | KLK13_HUMAN Kallikrein-13 OS=Homo sapiens OX=9606 GN=KLK13 PE=2 SV=1 | KLK13_HUMAN | -1.065249545 | 1.608010898 |
| 19 | Q9Y6K8;P00568 | KAD5_HUMAN Adenylate kinase isoenzyme 5 OS=Homo sapiens OX=9606 GN=AK5 PE=1 SV=2;KAD1_HUMAN Adenylate kinase isoenzyme 1 OS=Homo sapiens OX=9606 GN=AK1 PE=1 SV=3 | KAD5_HUMAN;KAD1_HUMAN | -1.068039205 | 2.576517843 |
| 20 | P80303 | NUCB2_HUMAN Nucleobindin-2 OS=Homo sapiens OX=9606 GN=NUCB2 PE=1 SV=3 | NUCB2_HUMAN | -1.079697223 | 1.605414899 |
| 21 | Q9UHG2 | PCS1N_HUMAN ProSAAS OS=Homo sapiens OX=9606 GN=PCSK1N PE=1 SV=1 | PCS1N_HUMAN | -1.081094387 | 1.805481752 |
| 22 | Q02539;P16402;P10412;P16401;P16403 | H11_HUMAN Histone H1.1 OS=Homo sapiens OX=9606 GN=H1-1 PE=1 SV=3;H13_HUMAN Histone H1.3 OS=Homo sapiens OX=9606 GN=H1-3 PE=1 SV=2;H14_HUMAN Histone H1.4 OS=Homo sapiens OX=9606 GN=H1-4 PE=1 SV=2;H15_HUMAN Histone H1.5 OS=Homo sapiens OX=9606 | H11_HUMAN;H13_HUMAN;H14_HUMAN;H15_HUMAN;H12_HUM | -1.083484519 | 1.965200449 |
| 23 | P13861;P31323 | KAP2_HUMAN cAMP-dependent protein kinase type II-alpha regulatory subunit OS=Homo sapiens OX=9606 GN=PRKAR2A PE=1 SV=2;KAP3_HUMAN cAMP-dependent protein kinase type II-beta regulatory subunit OS=Homo sapiens OX=9606 GN=PRKAR2B PE=1 SV=3 | KAP2_HUMAN;KAP3_HUMAN | -1.093212283 | 3.275373962 |
| 24 | P02751 | FINC_HUMAN Fibronectin OS=Homo sapiens OX=9606 GN=FN1 PE=1 SV=5 | FINC_HUMAN | -1.097815047 | 2.024980525 |
| 25 | Q9UN37;O75351;Q6PIW4 | VPS4A_HUMAN Vacuolar protein sorting-associated protein 4A OS=Homo sapiens OX=9606 GN=VPS4A PE=1 SV=1;VPS4B_HUMAN Vacuolar protein sorting-associated protein 4B OS=Homo sapiens OX=9606 GN=VPS4B PE=1 SV=2;FIGL1_HUMAN Fidgetin-like protein 1 OS=Homo sapiens OX= | VPS4A_HUMAN;VPS4B_HUMAN;FIGL1_HUMAN | -1.099503422 | 1.847046904 |
| 26 | B0FP48;E5RIL1 | UPK3L_HUMAN Uroplakin-3b-like protein 1 OS=Homo sapiens OX=9606 GN=UPK3BL1 PE=2 SV=1;UPKL2_HUMAN Uroplakin-3b-like protein 2 OS=Homo sapiens OX=9606 GN=UPK3BL2 PE=3 SV=2 | UPK3L_HUMAN;UPKL2_HUMAN | -1.134004804 | 3.002357709 |
| 27 | P19883 | FST_HUMAN Follistatin OS=Homo sapiens OX=9606 GN=FST PE=1 SV=2 | FST_HUMAN | -1.135667028 | 1.388503658 |
| 28 | Q96BQ1 | FAM3D_HUMAN Protein FAM3D OS=Homo sapiens OX=9606 GN=FAM3D PE=1 SV=1 | FAM3D_HUMAN | -1.15367043 | 2.808679943 |
| 29 | P31323 | KAP3_HUMAN cAMP-dependent protein kinase type II-beta regulatory subunit OS=Homo sapiens OX=9606 GN=PRKAR2B PE=1 SV=3 | KAP3_HUMAN | -1.223045445 | 2.16837603 |
| 30 | P82987 | ATL3_HUMAN ADAMTS-like protein 3 OS=Homo sapiens OX=9606 GN=ADAMTSL3 PE=1 SV=4 | ATL3_HUMAN | -1.255224231 | 2.514144187 |
| 31 | P06731 | CEAM5_HUMAN Carcinoembryonic antigen-related cell adhesion molecule 5 OS=Homo sapiens OX=9606 GN=CEACAM5 PE=1 SV=4 | CEAM5_HUMAN | -1.264172224 | 2.106423238 |
| 32 | Q02818 | NUCB1_HUMAN Nucleobindin-1 OS=Homo sapiens OX=9606 GN=NUCB1 PE=1 SV=4 | NUCB1_HUMAN | -1.332420247 | 1.71495615 |
| 33 | Q16352;P17661;P08670;P07197 | AINX_HUMAN Alpha-internexin OS=Homo sapiens OX=9606 GN=INA PE=1 SV=2;DESM_HUMAN Desmin OS=Homo sapiens OX=9606 GN=DES PE=1 SV=3;VIME_HUMAN Vimentin OS=Homo sapiens OX=9606 GN=VIM PE=1 SV=4;NFM_HUMAN Neurofilament medium polypeptide OS=Homo sap | AINX_HUMAN;DESM_HUMAN;VIME_HUMAN;NFM_HUMAN | -1.342582271 | 1.84490708 |
| 34 | P08670;P41219 | VIME_HUMAN Vimentin OS=Homo sapiens OX=9606 GN=VIM PE=1 SV=4;PERI_HUMAN Peripherin OS=Homo sapiens OX=9606 GN=PRPH PE=1 SV=2 | VIME_HUMAN;PERI_HUMAN | -1.485988392 | 1.624359067 |
| 35 | Q86U17 | SPA11_HUMAN Serpin A11 OS=Homo sapiens OX=9606 GN=SERPINA11 PE=2 SV=2 | SPA11_HUMAN | -1.508169481 | 1.801072351 |
| 36 | Q13421 | MSLN_HUMAN Mesothelin OS=Homo sapiens OX=9606 GN=MSLN PE=1 SV=2 | MSLN_HUMAN | -1.527664391 | 2.468760396 |
| 37 | Q9UHD0 | IL19_HUMAN Interleukin-19 OS=Homo sapiens OX=9606 GN=IL19 PE=1 SV=2 | IL19_HUMAN | -1.537958474 | 1.34227949 |
| 38 | P05387 | RLA2_HUMAN Large ribosomal subunit protein P2 OS=Homo sapiens OX=9606 GN=RPLP2 PE=1 SV=1 | RLA2_HUMAN | -1.542517054 | 1.533589213 |
| 39 | P06731;P31997;P13688 | CEAM5_HUMAN Carcinoembryonic antigen-related cell adhesion molecule 5 OS=Homo sapiens OX=9606 GN=CEACAM5 PE=1 SV=4;CEAM8_HUMAN Carcinoembryonic antigen-related cell adhesion molecule 8 OS=Homo sapiens OX=9606 GN=CEACAM8 PE=1 SV=2;CEAM1_HUMAN Carcinoem | CEAM5_HUMAN;CEAM8_HUMAN;CEAM1_HUMAN | -1.698933797 | 1.62984066 |
| 40 | Q8TD84 | DSCL1_HUMAN Cell adhesion molecule DSCAML1 OS=Homo sapiens OX=9606 GN=DSCAML1 PE=1 SV=2 | DSCL1_HUMAN | -1.702831316 | 1.614502292 |
| 41 | P12109 | CO6A1_HUMAN Collagen alpha-1(VI) chain OS=Homo sapiens OX=9606 GN=COL6A1 PE=1 SV=3 | CO6A1_HUMAN | -1.749952243 | 2.044784786 |
| 42 | Q06828 | FMOD_HUMAN Fibromodulin OS=Homo sapiens OX=9606 GN=FMOD PE=1 SV=2 | FMOD_HUMAN | -1.987337021 | 1.612398563 |
| 43 | Q6ZMB0 | B3GN6_HUMAN Acetylgalactosaminyl-O-glycosyl-glycoprotein beta-1,3-N-acetylglucosaminyltransferase OS=Homo sapiens OX=9606 GN=B3GNT6 PE=1 SV=2 | B3GN6_HUMAN | -2.009863983 | 2.523581903 |
| 44 | P02452 | CO1A1_HUMAN Collagen alpha-1(I) chain OS=Homo sapiens OX=9606 GN=COL1A1 PE=1 SV=6 | CO1A1_HUMAN | -2.116755067 | 1.634518276 |
| 45 | Q8N4F0 | BPIB2_HUMAN BPI fold-containing family B member 2 OS=Homo sapiens OX=9606 GN=BPIFB2 PE=1 SV=2 | BPIB2_HUMAN | -2.348867328 | 2.088723077 |
| 46 | P08123 | CO1A2_HUMAN Collagen alpha-2(I) chain OS=Homo sapiens OX=9606 GN=COL1A2 PE=1 SV=7 | CO1A2_HUMAN | -2.492380367 | 1.613884262 |
| 47 | P13611 | CSPG2_HUMAN Versican core protein OS=Homo sapiens OX=9606 GN=VCAN PE=1 SV=3 | CSPG2_HUMAN | -2.540201539 | 3.002810119 |
| 48 | P08572 | CO4A2_HUMAN Collagen alpha-2(IV) chain OS=Homo sapiens OX=9606 GN=COL4A2 PE=1 SV=4 | CO4A2_HUMAN | -3.090663207 | 1.386387453 |
| 49 | P08118 | MSMB_HUMAN Beta-microseminoprotein OS=Homo sapiens OX=9606 GN=MSMB PE=1 SV=1 | MSMB_HUMAN | -4.169583731 | 1.688535753 |

| | A | B | C | D | E |
|---|---|---|---|---|---|
| 1 | Protein | Protein.Description | Protein.Name | WT-MUC5BKO_log2FC | NEG LOG 10 (PVALUE) |
| 2 | P08263 | GSTA1_HUMAN Glutathione S-transferase A1 OS=Homo sapiens OX=9606 GN=GSTA1 PE=1 SV=3 | GSTA1_HUMAN | 1.543070743 | 2.055084677 |
| 3 | Q9Y4F9 | RIPR2_HUMAN Rho family-interacting cell polarization regulator 2 OS=Homo sapiens OX=9606 GN=RIPOR2 PE=1 SV=4 | RIPR2_HUMAN | 1.310123843 | 2.257644609 |
| 4 | A6NCS6 | CB072_HUMAN Uncharacterized protein C2orf72 OS=Homo sapiens OX=9606 GN=C2orf72 PE=1 SV=2 | CB072_HUMAN | 1.287050784 | 2.159374198 |
| 5 | P05155 | IC1_HUMAN Plasma protease C1 inhibitor OS=Homo sapiens OX=9606 GN=SERPING1 PE=1 SV=2 | IC1_HUMAN | 1.232807747 | 1.788029346 |
| 6 | O43927 | CXL13_HUMAN C-X-C motif chemokine 13 OS=Homo sapiens OX=9606 GN=CXCL13 PE=1 SV=1 | CXL13_HUMAN | 1.213137708 | 1.697838344 |
| 7 | O75190 | DNJB6_HUMAN DnaJ homolog subfamily B member 6 OS=Homo sapiens OX=9606 GN=DNAJB6 PE=1 SV=2 | DNJB6_HUMAN | 1.203766861 | 2.243111378 |
| 8 | A6QL63 | ABTB3_HUMAN Ankyrin repeat and BTB/POZ domain-containing protein 3 OS=Homo sapiens OX=9606 GN=ABTB3 PE=2 SV=3 | ABTB3_HUMAN | 1.04755224 | 1.487770852 |
| 9 | P25311 | ZA2G_HUMAN Zinc-alpha-2-glycoprotein OS=Homo sapiens OX=9606 GN=AZGP1 PE=1 SV=2 | ZA2G_HUMAN | 1.022791563 | 2.055890315 |
| 10 | Q9H3R2 | MUC13_HUMAN Mucin-13 OS=Homo sapiens OX=9606 GN=MUC13 PE=1 SV=3 | MUC13_HUMAN | 1.003451225 | 1.407021702 |
| 11 | P19883 | FST_HUMAN Follistatin OS=Homo sapiens OX=9606 GN=FST PE=1 SV=2 | FST_HUMAN | -1.017806207 | 1.341063794 |
| 12 | Q14210 | LY6D_HUMAN Lymphocyte antigen 6D OS=Homo sapiens OX=9606 GN=LY6D PE=1 SV=1 | LY6D_HUMAN | -1.043105806 | 1.339981825 |
| 13 | P58876 | H2B1D_HUMAN Histone H2B type 1-D OS=Homo sapiens OX=9606 GN=H2BC5 PE=1 SV=2 | H2B1D_HUMAN | -1.09108579 | 1.326303514 |
| 14 | P16402;P10412;P16403 | H13_HUMAN Histone H1.3 OS=Homo sapiens OX=9606 GN=H1-3 PE=1 SV=2;H14_HUMAN Histone H1.4 OS=Homo sapiens OX=9606 GN=H1-4 PE=1 SV=2;H12_HUMAN Histone H1.2 OS=Homo sapiens OX=9606 GN=H1-2 PE=1 SV=2 | H13_HUMAN;H14_HUMAN;H12_HUMAN | -1.09881781 | 1.594043512 |
| 15 | P13473 | LAMP2_HUMAN Lysosome-associated membrane glycoprotein 2 OS=Homo sapiens OX=9606 GN=LAMP2 PE=1 SV=2 | LAMP2_HUMAN | -1.110730316 | 1.40235207 |
| 16 | Q5QNW6;P23527;Q93079 | H2B2F_HUMAN Histone H2B type 2-F OS=Homo sapiens OX=9606 GN=H2BC18 PE=1 SV=3;H2B1O_HUMAN Histone H2B type 1-O OS=Homo sapiens OX=9606 GN=H2BC17 PE=1 SV=3;H2B1H_HUMAN Histone H2B type 1-H OS=Homo sapiens OX=9606 GN=H2BC9 PE=1 SV=3 | H2B2F_HUMAN;H2B1O_HUMAN;H2B1H_HUMAN | -1.147127149 | 1.390318118 |
| 17 | P05387 | RLA2_HUMAN Large ribosomal subunit protein P2 OS=Homo sapiens OX=9606 GN=RPLP2 PE=1 SV=1 | RLA2_HUMAN | -1.180578664 | 1.314427377 |
| 18 | O43252 | PAPS1_HUMAN Bifunctional 3'-phosphoadenosine 5'-phosphosulfate synthase 1 OS=Homo sapiens OX=9606 GN=PAPSS1 PE=1 SV=2 | PAPS1_HUMAN | -1.199161932 | 1.37727948 |
| 19 | P17405 | ASM_HUMAN Sphingomyelin phosphodiesterase OS=Homo sapiens OX=9606 GN=SMPD1 PE=1 SV=5 | ASM_HUMAN | -1.218736734 | 1.962041478 |
| 20 | P13611 | CSPG2_HUMAN Versican core protein OS=Homo sapiens OX=9606 GN=VCAN PE=1 SV=3 | CSPG2_HUMAN | -1.242464105 | 2.692226916 |
| 21 | P16401 | H15_HUMAN Histone H1.5 OS=Homo sapiens OX=9606 GN=H1-5 PE=1 SV=3 | H15_HUMAN | -1.244730696 | 2.188062359 |
| 22 | Q02818 | NUCB1_HUMAN Nucleobindin-1 OS=Homo sapiens OX=9606 GN=NUCB1 PE=1 SV=4 | NUCB1_HUMAN | -1.266242166 | 1.67204303 |
| 23 | Q02539;P16402;P10412;P16401;P16403 | H11_HUMAN Histone H1.1 OS=Homo sapiens OX=9606 GN=H1-1 PE=1 SV=3;H13_HUMAN Histone H1.3 OS=Homo sapiens OX=9606 GN=H1-3 PE=1 SV=2;H14_HUMAN Histone H1.4 OS=Homo sapiens OX=9606 GN=H1-4 PE=1 SV=2;H15_HUMAN Histone H1.5 OS=Homo sapiens OX=9606 GN=H1-5 PE=1 SV=3;H12_HUMAN Histone H1.2 OS=H | H11_HUMAN;H13_HUMAN;H14_HUMAN;H15_HUMAN;H12_HUMAN | -1.314243414 | 2.130646749 |
| 24 | Q9UHG2 | PCS1N_HUMAN ProSAAS OS=Homo sapiens OX=9606 GN=PCSK1N PE=1 SV=1 | PCS1N_HUMAN | -1.462926053 | 2.06356595 |
| 25 | P98088;Q02817;Q9HC84 | MUC5A_HUMAN Mucin-5AC OS=Homo sapiens OX=9606 GN=MUC5AC PE=1 SV=4;MUC2_HUMAN Mucin-2 OS=Homo sapiens OX=9606 GN=MUC2 PE=1 SV=3;MUC5B_HUMAN Mucin-5B OS=Homo sapiens OX=9606 GN=MUC5B PE=1 SV=3 | MUC5A_HUMAN;MUC2_HUMAN;MUC5B_HUMAN | -1.508338582 | 1.753453371 |
| 26 | P98088 | MUC5A_HUMAN Mucin-5AC OS=Homo sapiens OX=9606 GN=MUC5AC PE=1 SV=4 | MUC5A_HUMAN | -1.515835651 | 2.73466488 |
| 27 | Q6ZMB0 | B3GN6_HUMAN Acetylgalactosaminyl-O-glycosyl-glycoprotein beta-1,3-N-acetylglucosaminyltransferase OS=Homo sapiens OX=9606 GN=B3GNT6 PE=1 SV=2 | B3GN6_HUMAN | -1.696883378 | 2.377337048 |
| 28 | Q86U17 | SPA11_HUMAN Serpin A11 OS=Homo sapiens OX=9606 GN=SERPINA11 PE=2 SV=2 | SPA11_HUMAN | -1.821608426 | 1.961845135 |
| 29 | Q06828 | FMOD_HUMAN Fibromodulin OS=Homo sapiens OX=9606 GN=FMOD PE=1 SV=2 | FMOD_HUMAN | -1.890853288 | 1.590807143 |
| 30 | Q8N4F0 | BPIB2_HUMAN BPI fold-containing family B member 2 OS=Homo sapiens OX=9606 GN=BPIFB2 PE=1 SV=2 | BPIB2_HUMAN | -2.028012416 | 1.962951176 |
| 31 | P08118 | MSMB_HUMAN Beta-microseminoprotein OS=Homo sapiens OX=9606 GN=MSMB PE=1 SV=1 | MSMB_HUMAN | -3.569486251 | 1.558360633 |
